# A mutation in the endonuclease domain of mouse MLH3 reveals novel roles for MutLγ during crossover formation in meiotic prophase I

**DOI:** 10.1101/517748

**Authors:** Melissa Toledo, Xianfei Sun, Miguel A. Brieño-Enríquez, Vandana Raghavan, Stephen Gray, Jeffrey Pea, Anita Venkatesh, Lekha Patel, Peter L. Borst, Eric Alani, Paula E. Cohen

## Abstract

During meiotic prophase I, double-strand breaks (DSBs) initiate homologous recombination leading to non-crossovers (NCOs) and crossovers (COs). In mouse, 10% of DSBs are designated to become COs, primarily through a pathway dependent on the MLH1-MLH3 heterodimer (MutLγ). Mlh3 contains an endonuclease domain that is critical for resolving COs in yeast. We generated a mouse (*Mlh3^DN/DN^*) harboring a mutation within this conserved domain that is predicted to generate a protein that is catalytically inert. *Mlh3^DN/DN^* males, like fully null *Mlh3^-/-^* males, have no spermatozoa and are infertile, yet spermatocytes have normal DSBs and undergo normal synapsis events in early prophase I. Unlike *Mlh3^-/-^* males, mutation of the endonuclease domain within MLH3 permits normal loading and frequency of MutLγ in pachynema. However, key DSB repair factors (RAD51) and mediators of CO pathway choice (BLM helicase) persist into pachynema in *Mlh3^DN/DN^* males, indicating a temporal delay in repair events and revealing a mechanism by which alternative DSB repair pathways may be selected. While *Mlh3^DN/DN^* spermatocytes retain only 22% of wildtype chiasmata counts, this frequency is greater than observed in *Mlh3^-/-^* males (10%), suggesting that the allele may permit partial endonuclease activity, or that other pathways can generate COs from these MutLγ-defined repair intermediates in *Mlh3^DN/DN^* males. Double mutant mice homozygous for the *Mlh3^DN/DN^* and *Mus81^-/-^* mutations show losses in chiasmata that approach levels observed in *Mlh3^-/-^* males, indicating that the MUS81-EME1-regulated crossover pathway accounts for some of the increased residual chiasmata observed in *Mlh3^DN/DN^* spermatocytes. Our data demonstrate that mouse spermatocytes bearing the MLH1-MLH3^DN/DN^ complex display the proper loading of factors essential for CO resolution (MutSγ, CDK2, HEI10, MutLγ). Despite these functions, mice bearing the *Mlh3^DN/DN^* allele show defects in the repair of meiotic recombination intermediates and a loss of most chiasmata.

**SUMMARY:** The MLH1-MLH3 complex is essential for crossing over in mammalian meiosis. We generated a mutation in mouse MLH3 that alters its conserved endonuclease domain and show that it disrupts crossing over in a manner distinct from the full null *Mlh3* mouse, but also results in male infertility.

## INTRODUCTION

Meiosis is a specialized cell division process in which a diploid parental cell undergoes one round of DNA replication followed by two rounds of division, resulting in up to four haploid gametes. Successful halving of the genome during meiosis I depends on the tethering of maternal and paternal homologous chromosomes during meiotic prophase I, and their subsequent release at the first meiotic division. This tethering is ensured by homologous recombination, leading to the formation of crossovers; by synapsis, the formation of a tripartite proteinaceous structure, the synaptonemal complex, or SC between homologous chromosomes; and by cohesion between replicated sister chromatids that ensures appropriate tension on the metaphase I spindle (Gray and Cohen 2016; Hunter 2015). Thus, recombination and synapsis are hallmarks of prophase I, and are both essential for ensuring homolog interactions leading to the formation of at least one crossover event per chromosome pair. Moreover, the correct placement, frequency, and distribution of crossovers is critical for ensuring appropriate disjunction at metaphase I and for maintaining genomic stability (Gray and Cohen 2016; Bolcun-Filas and Handel 2018).

Meiotic recombination begins with the introduction of a large number of programmed double-strand breaks (DSBs), which are repaired as non-crossovers (NCOs) or crossovers (COs). Evidence for distinct NCO versus CO pathways was obtained in *S. cerevisiae*, where it was shown that the former occur earlier in meiotic prophase I, and subsequent work suggested that they appeared primarily through synthesis-dependent strand annealing (SDSA)(McMahill et al. 2007; Allers and Lichten 2001). In *M. musculus*, 90% of DSBs are repaired as NCOs, presumably via SDSA or other pathways, while 10% of DSBs are repaired as COs (Holloway et al. 2008; Guillon et al. 2005; Svetlanov et al. 2008; Cole et al. 2010).

COs can form via one of at least two distinct mechanisms (referred to as class I and class II), each of which is used in varying degrees in different eukaryotic organisms (de los Santos et al. 2003; Higgins et al. 2008; Holloway et al. 2008). The class I CO pathway is also known as the ZMM pathway, named after the major genes discovered in yeast that regulate this mechanism (Lynn et al. 2007; Jessop et al. 2006; Börner et al. 2004; Wang et al. 2009; Agarwal and Roeder 2000; Hollingsworth et al. 1995; Tsubouchi et al. 2006). Class II COs, on the other hand, do not involve the ZMM proteins, but instead appear to rely on the structure-specific endonuclease (SSN), MUS81/EME1 (Mus81/Mms4 in *S. cerevisiae*) (de los Santos et al. 2003; Higgins et al. 2008; Holloway et al. 2008). Class I COs differ from class II COs in that the former are regulated by interference, the process by which placement of one CO prevents the nearby localization of a second CO, thus resulting in appropriate spacing of these events across the genome (de Boer et al. 2006).

In the class I CO pathway, DSBs are processed and resected to form single-end invasion (SEI) intermediates. This is followed by displacement of single strand DNA (ssDNA) from the recipient homolog to produce a double Holliday junction (dHJ). The yeast ZMM proteins Msh4 and Msh5 form a complex known as MutSγ that associates with a subset of these intermediate structures (Novak et al. 2001; Nishant et al. 2010; Pochart et al. 1997). At least in yeast, this recruitment may be dependent on the STR complex, consisting of Sgs1 (BLM in mammals), <u>T</u>op3 and <u>R</u>mi1 (Kaur et al. 2015; Tang et al. 2015). STR is proposed to act by disassembling the early recombination intermediates that would otherwise be processed through SSN-directed recombination pathways, thereby promoting either early NCO formation via SDSA, or CO formation through the capture of these recombination intermediates by the ZMM proteins, including MutSγ (Kaur et al. 2015). MutSγ is then thought to stabilize the dHJs, leading to the recruitment of a second MMR complex, MutLγ, consisting of the MutL homologs, Mlh1 and Mlh3 (Hunter and Borts 1997; Wang et al. 1999). The mouse MutSγ complex associates with chromosome cores in zygonema (Kneitz et al. 2000), recruiting the MutLγ complex in pachynema. However, MutLγ associates with only a subset of MutSγ sites (~24-26 and 150 foci/nucleus, respectively), designating these events as class I COs (Kolas et al. 2005; Lipkin et al. 2002).

Though not formerly considered to be ZMM proteins, MLH1 and MLH3 are critical for class I CO events in numerous organisms (Al-Sweel et al. 2017; Woods et al. 1999; Wang et al. 1999; Lipkin et al. 2002; Nishant et al. 2008; Argueso et al. 2002; Rogacheva et al. 2014). In fact, the *M. musculus* MLH1-MLH3 heterodimer localizes to sites that are destined to become class I COs and the absence of either subunit in male spermatocytes leads to a dramatic decrease, but not complete absence, of chiasmata (the physical manifestation of a CO) (Kan et al. 2008; Kolas et al. 2005; Marcon and Moens 2003; Lipkin et al. 2002; Baker et al. 1996; Anderson et al. 1999). While MutLγ is known to be recruited to sites that are preloaded with MutSγ, recent studies have shown that *S. cerevisiae* MutLγ can bind to single and double-stranded DNA (ssDNA, dsDNA), as well as a variety of branched DNA structures (Rogacheva et al. 2014; Manhart et al. 2017; Claeys Bouuaert and Keeney 2017; Ranjha et al. 2014). How such binding properties relate to the *in vivo* functions of MutLγ remains unclear.

Class I CO formation in *M. musculus* is dependent on MLH3, and on its heterodimeric interaction with MLH1 (Lipkin et al. 2002; Svetlanov et al. 2008; Kan et al. 2008). Interestingly, MLH3 recruitment precedes that of MLH1 (Kolas et al. 2005). Further analysis of MutLγ has shown that MLH3 contains a conserved metal binding motif, DQHA(X)_2_E(X)_4_E, originally discovered in the human MutL homolog, PMS2, and found to be required for human MutLα (hMLH1/hPMS2) endonuclease function (Kadyrov et al. 2006). This putative endonuclease motif is highly conserved in eukaryotic homologs of human PMS2 and MLH3, but not in homologs of human MLH1 and PMS1. The expectation for MLH3 is that this endonuclease function might represent a “resolvase” activity for class I COs. Studies in *S. cerevisiae* have shown that a single point mutation in the endonuclease motif of yeast Mlh3 (*mlh3-D523N*) disrupts its endonucleolytic activity and results in meiotic crossover defects similar to full *mlh3* (*mlh3Δ*) null mutants, yet does not affect the protein stability of Mlh3 or its interaction with Mlh1 (Nishant et al. 2008). Further analysis of the entire endonuclease domain in *S. cerevisiae* revealed that mutation of any conserved residue results in a null or near-null phenotype with respect to crossing over (Al-Sweel et al. 2017). Biochemical analysis reveals that the Mlh1-mlh3D523N protein lacks the ability to nick closed circular double stranded DNA, indicating loss of endonuclease activity (Rogacheva et al. 2014; Ranjha et al. 2014). Collectively, these studies in *S. cerevisiae* suggest that MutLγ plays a direct role in resolving dHJs to generate COs through its endonuclease activity.

To investigate the function of the putative endonuclease domain of MLH3 in mammalian meiotic recombination, we generated a point mutant mouse (termed *Mlh3^DN^*) in which the endonucleolytic domain was disrupted at the orthologous residue to the D523N mutation in yeast, allowing the overall structure of MLH3 to remain intact, as determined by the ability to form a stable complex with MLH1. By generating a catalytically defective protein, we hypothesized that the mutant MutLγ complex would remain structurally intact and thus might reveal a functional interplay with other meiotic CO functions. We demonstrate that normal function of the MLH3 endonuclease domain is required for resolution of DSB repair intermediates towards CO formation and thus for late meiotic recombination events. *Mlh3^DN/DN^* spermatocytes exhibit normal DSB formation and early processing, and normal synapsis through early prophase I. *Mlh3^DN/DN^* spermatocytes exhibit appropriate localization of MLH3 and MLH1 to the synaptonemal complex during pachynema, along with pro-crossover factors HEI10 and CDK2, phenotypes that are clearly different from that observed in *Mlh3^−/−^* males. However, *Mlh3^DN/DN^* diakinesis-staged spermatocytes show significantly fewer chiasmata compared to wild-type mice (WT), but significantly more when compared to *Mlh3^−/−^* males, suggesting either that the MLH3^DN^ protein retains partial endonucleolytic activity, or that the presence of the MutLγ complex, albeit altered in its endonucleolytic capacity, can invoke other repair pathways to become active. Interestingly, in line with the latter suggestion, we find that the RecQ helicase, BLM, is upregulated throughout prophase I in *Mlh3^DN/DN^* spermatocytes, perhaps aiding the recruitment of other repair proteins. To explore the increase in residual chiasmata observed at diakinesis in *Mlh3^DN/DN^* males relative to that of *Mlh3^−/−^* males, we demonstrate that co-incident loss of the class II CO pathway in *Mus81^−/−^Mlh3^DN/DN^* double mutant males results in altered distribution of MutLγ, with an increased proportion of synapsed autosomes bearing no MutLγ foci. Furthermore, the proportion of chiasmata remaining in these double mutants is between that of *Mlh3^DN/DN^* and *Mlh3^−/−^* males, suggesting that MUS81-EME1 accounts for a significant proportion of these additional chiasmata. Collectively, our data show that the endonucleolytic activity of MLH3 is important for normal processing of DSB repair intermediates through the Class I pathway.

## RESULTS

### *Mlh3^DN/DN^* males are infertile

To investigate the meiotic requirement for the presence of a functional endonuclease domain in mammalian MLH3, we generated a mouse line with a point mutation in a conserved endonuclease motif located in the *M. musculus* protein: DQHAAHERIRLE (Gueneau et al., 2013; Kadyrov et al., 2006). Specifically, we replaced the aspartic acid “D” in amino acid position 1185, with an asparagine "N" by changing GAC to AAC in the genomic sequence, termed MLH3^DN^ throughout. Extrapolating from an analogous mutation in the *S. cerevisiae* gene, this D-to-N replacement is predicted to disrupt the endonucleolytic function of MLH3 while maintaining the ability to interact stably with MLH1 ((Nishant et al., 2008); Figure S1). Mice were maintained on a C57Bl/6J background throughout the study.

Male *Mlh3^+/DN^* mice were phenotypically similar to WT littermates and displayed full fertility. *Mlh3^DN/DN^* males are also grossly normal when compared to WT littermates, survive into adulthood, and live normal lifespans. *Mlh3^DN/DN^* males also exhibit normal mating behaviors as determined by observing a vaginal plug in WT females the morning after mating. However, breeding between multiple sets of *Mlh3^DN/DN^* males and WT females was never observed to result in offspring over a four-year period.

Similar to the situation seen for *Mlh3^−/−^* males (Lipkin et al., 2002), *Mlh3^DN/DN^* males show complete infertility, accompanied by significantly reduced testes size when compared to WT (Figure 1A, B; p < 0.0001) and the absence of spermatozoa in the epididymides (Figure 1C; p < 0.0001). Whereas histological cross-sections of testes stained with hemotoxylin and eosin from WT males showed the presence of meiotic and post-meiotic cells within the seminiferous epithelium, testis sections from *Mlh3^DN/DN^* males were devoid of spermatids, but showed the presence of spermatogonia and spermatocytes (Figure 1D-G). In addition, metaphase I spermatocytes were observed in the tubular lumen of *Mlh3^DN/DN^* mice (Figure 1G, black arrows). Thus, mutation of the endonuclease domain of *Mlh3* in the mouse results in a sterility phenotype grossly similar to that seen in *Mlh3^−/−^* mice.

**Figure 1.**
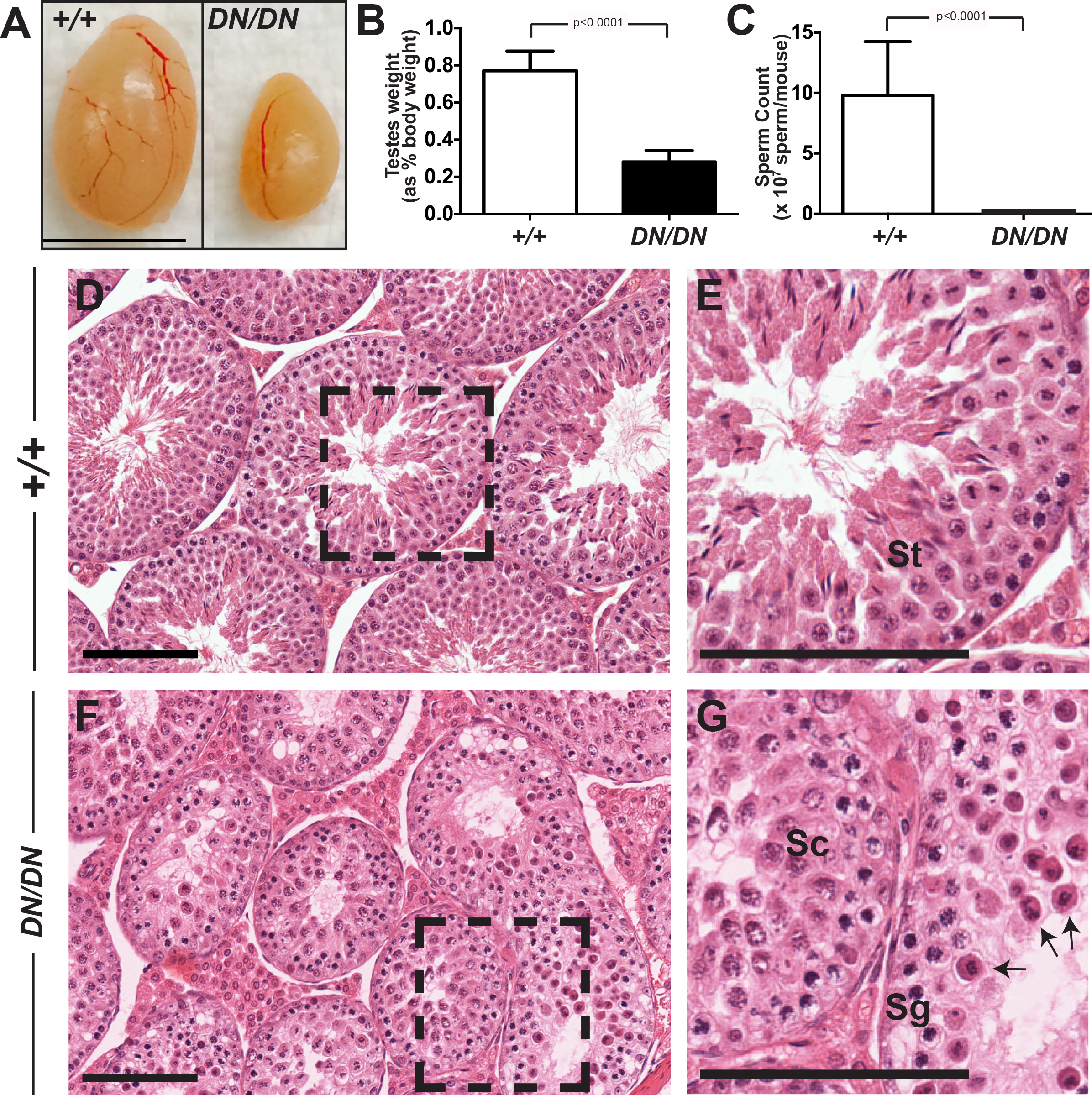
*Mlh3^DN/DN^* males show a sterile phenotype. (A, B) *Mlh3^DN/DN^* adult male testes are significantly smaller when compared to WT littermates (*Mlh3^DN/DN^* - 0.28% of total body weight ± 0.06, n = 13; WT - 0.77% ± 0.1, n = 15; p < 0.0001, unpaired t-test with Welch's correction) and (C) have zero sperm in the epididymis (WT - 9.8 × 10^7^ ± 4.4 sperm/mouse; p < 0.0001; unpaired t-test with Welch's correction, n = 10 and 12 mice, respectively; error bars show standard deviation). Hemotoxylin and eosin stained (D, E) WT testes show the presence of meiotic and post-meiotic cells whereas (F, G) *Mlh3^DN/DN^* are absent of spermatids and spermatozoa. Higher magnification of WT and *Mlh3^DN/DN^* testes sections are shown in E and G, respectively. Black arrows in (G) indicate metaphase I spermatocytes in the *Mlh3^DN/DN^*seminiferous tubule lumen. Sg, spermatogonia; Sc, prophase I spermatocytes; St, postmeiotic spermatids. Scale bar is 100 μm. Images were taken at 40X magnification.

### *Mlh3*^*DN/DN*^ spermatocytes, like those of *Mlh3*^*−/−*^ mice, exhibit normal DSB formation and synapsis

To investigate the progression of meiotic recombination, prophase I chromosome spreads were prepared from WT, *Mlh3^DN/DN^*, and *Mlh3^−/−^* adult males and stained for a variety of markers involved in synapsis and recombination. Chromosome spreads were stained with antibodies against γH2AX, the phosphorylated form of histone H2AX, as a marker of DSBs (Viera et al. 2004; Mahadevaiah et al. 2001). In spermatocyte preparations from WT males, γH2AX signal is abundant throughout the nucleus at leptonema, coincident with the induction of several hundred DSBs (Mahadevaiah et al. 2001; Gray and Cohen 2016). The γH2AX signal declines in zygonema as DSBs are processed for repair (Mahadevaiah et al. 2001; Hunter et al. 2001). In pachynema and diplonema, γH2AX signal is absent from the autosomes, but emerges throughout the sex body due to meiotic sex chromosome inactivation (MSCI) ((Handel 2004); Figure S2A-D). Spermatocytes from both *Mlh3^DN/DN^* and *Mlh3^−/−^* males exhibit the same γH2AX signal and temporal dynamics as observed in WT spermatocytes, with abundant staining in leptonema, slightly reduced signaling in zygonema, followed by the absence of γH2AX signal on the autosomes of pachytene and diplotene spermatocytes, except at the sex body (Figure S2F-I, K-N). We do not see specific persistent γH2AX signal on the autosomes at pachynema in *Mlh3^−/−^* spermatocytes (Qiao et al. 2014), unless we markedly increase our imaging exposure time γH2AX (Figure S2E, J, O; white arrows). Under these conditions, we see persistent foci of γH2AX in spermatocytes from WT and from *Mlh3^DN/DN^* spermatocytes also. Thus, in our hands, we see no specific persistence in autosomal γH2AX signal through pachynema in mice lacking MLH3 or harboring a mutation within the endonuclease domain of MLH3.

Spermatocyte chromosome spreads from WT and *Mlh3^DN/DN^* males were stained with antibodies against synaptonemal complex (SC) components, SYCP3 and SYCP1, marking the axial/lateral elements and the central element, respectively. Prophase I progression in WT spreads is characterized by the initial accumulation of SYCP3 signal in discrete dots along chromosomes at leptonema, and these dots gradually coalesce into continuous filaments along the chromosome cores in zygonema (Figure S2Q). At this time, SYCP1 appears in patches along the SYCP3 signal, indicating that synapsis is occurring. By late zygonema, most of the chromosome core is now labeled with SYCP1, and by pachynema synapsis is complete, as demonstrated by complete overlap of the SYCP3/SYCP1 signals on the autosomes. For the sex chromosomes, synapsis only occurs at the pseudoautosomal region (PAR). After meiotic recombination occurs, the SC begins to degrade in diplonema, and the homologs are no longer tethered to one another except at CO sites (Figure S2P-S).

Synapsis appears normal in *Mlh3^DN/DN^* spermatocytes with discrete accumulation of SYCP3 on the chromosomes in leptonema, followed by continued accumulation of SYCP3 along the chromosomes as SYCP1 appears in patches in zygonema (Figure S2T, U). Complete synapsis of the autosomes and the PAR is observed in pachynema with co-localization of SYCP1 and SYCP3 (Figure S2V). Desynapsis is then observed in diplonema with the degradation of the SC (Figure S2W). Thus, synapsis in *Mlh3^DN/DN^* spermatocytes appears unaffected by loss of the endonuclease activity of MLH3, a result similar to that seen for complete loss of MLH3 protein.

### *Mlh3^DN/DN^* spermatocytes show a persistence of RAD51 in pachytene cells

Early DSB repair events were monitored by examining and quantifying localization of the RecA strand exchange protein, RAD51, on chromosome cores of the autosomes throughout prophase I (Ashley et al. 1995). In WT mice, RAD51 localizes to chromosome cores of early and late zygotene cells as discrete foci at a high frequency (EZ and LZ, respectively; Figure 2A, G). Compared to WT littermates in early and late zygonema, RAD51 counts in spermatocytes from *Mlh3^DN/DN^* males were significantly elevated (Figure 2C,G; p<0.001 and p<0.01, respectively, by unpaired t-test with Welch’s correction). However, while early zygotene RAD51 counts were indistinguishable in *Mlh3^−/−^* spermatocytes compared to WT (Figure 2E,G), they were significantly lower than that seen at the equivalent stage in *Mlh3^DN/DN^* males (p<0.001 by unpaired t-test with Welch’s correction). By late zygonema, the RAD51 counts in were significantly lower in *Mlh3^−/−^* spermatocytes compared to WT and *Mlh3^DN/DN^* animals (p<0.001 by unpaired t-test with Welch’s correction).

**Figure 2.**
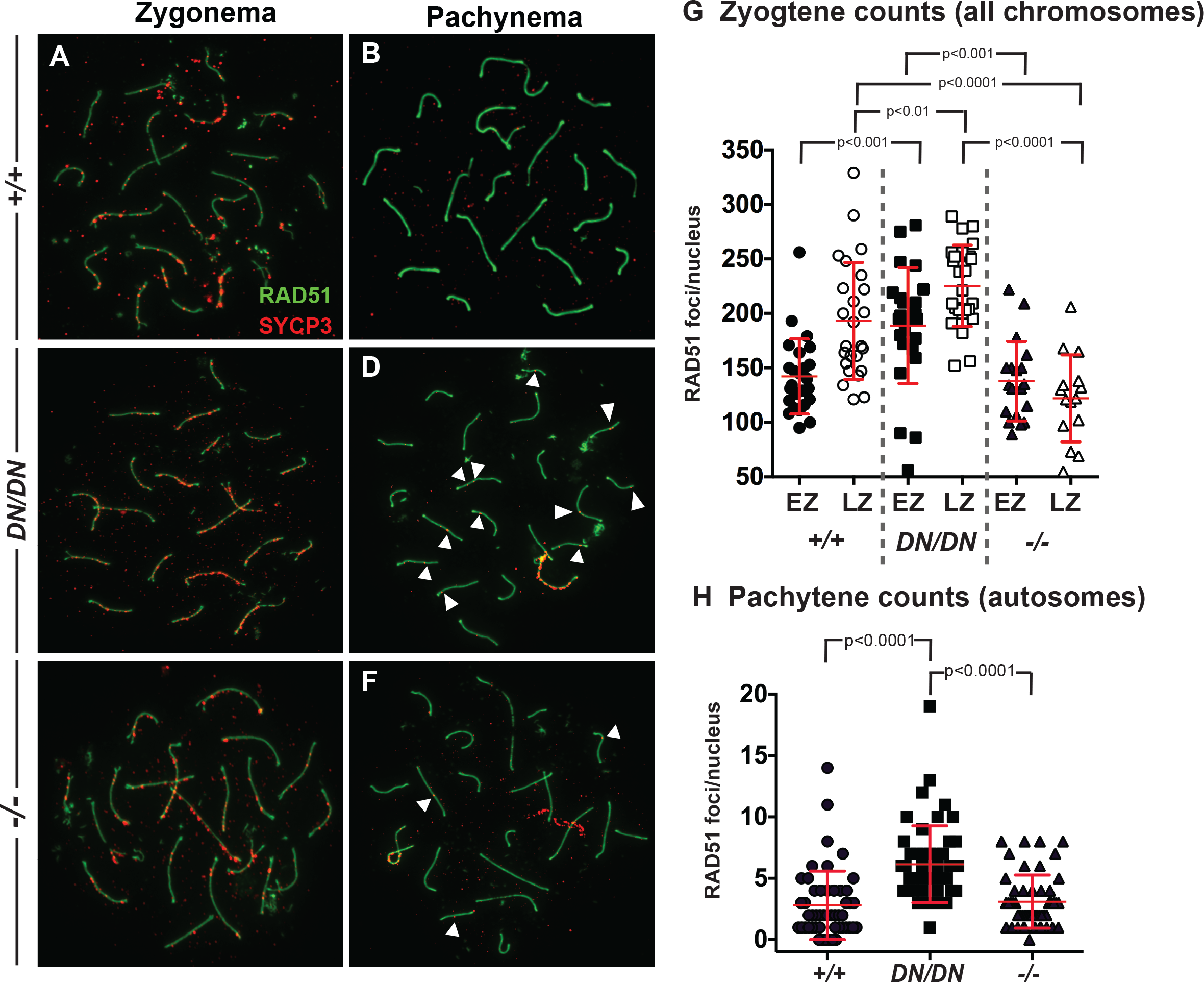
*Mlh3^DN/DN^* spermatocytes show a persistence of RAD51 foci in pachynema. The localization of RAD51 (green) on synaptonemal complex protein SYCP3 (red) is observed in (A, B) WT and (C, D) *Mlh3^DN/DN^* male spermatocytes in zygonema and pachynema. (A, C, E, G) In early zygonema, WT cells show high numbers of RAD51 associated with the chromosome cores while *Mlh3^DN/DN^* exhibit even higher RAD51 foci (WT mean + standard deviation = 142.1 ± 34.5 foci, *Mlh3^DN/DN^* mean + standard deviation = 188.8 ± 53.3 foci, p<0.001 by unpaired t-test with Welch's correction). In late zygonema, *Mlh3^DN/DN^*spermatocytes continue to have higher RAD51 foci than WT (WT mean + standard deviation = 193.0 ± 53.7 foci, *Mlh3^DN/DN^* mean ± standard deviation = 225.3 ± 37.3 foci; p<0.01 by unpaired t-test with Welch’s correction). *Mlh3^−/−^* late zygotene cells show significantly fewer RAD51 focus counts (mean = 122.1 ± 39.9 foci) when compared to WT and *Mlh3^DN/DN^* (both p < 0.0001 by unpaired t-test with Welch’s correction). In all cases, at least 3 mice were assessed for each genotype and at least 20 cells per mouse (B, D, F, H) In pachynema, WT cells exhibit a dramatic decrease of very few to no RAD51 associated with the autosomes while *Mlh3^DN/DN^* cells show persistent RAD51 (WT mean ± standard deviation = 2.8 ± 2.8 foci, *Mlh3^DN/DN^* mean ± standard deviation = 6.1 ± 3.1 foci; p<0.0001, unpaired t-test). *Mlh3^−/−^* pachytene cells exhibit comparable RAD51 focus counts (mean + standard deviation = 3.1 ± 2.1 foci) when compared to WT (p = 0.55 by unpaired t-test), but significantly fewer than *Mlh3^DN/DN^* spermatocytes(p<0.0001 by unpaired t-test). Note that sex chromosome-associated RAD51 staining was excluded from counts at pachynema. For all chromosome imaging and foci counts, at least three mice of each genotype were observed for each staining set.

By pachynema, RAD51 foci frequency in spermatocytes from WT decreased to very low numbers, as did that of *Mlh3^−/−^* males (Figures 2B,F,H; p=0.55 unpaired t-test). In contrast, focus counts in pachytene spermatocytes from *Mlh3^DN/DN^* males were significantly elevated (Figure 2D,H; p<0.0001). These observations suggest a persistence of DSB repair intermediates loaded with RecA homologs in *Mlh3^DN/DN^* spermatocytes in pachynema, or an elongated window of DSB induction in these mutant animals. We hypothesize, based on these observations, that RAD51 accumulation in zygonema is affected by loss or mutation of MLH3 protein, suggesting an early function for MutLγ in establishing appropriate DSB repair intermediates that is not confined to CO pathway fate. Further, we hypothesize that DSBs are repaired in a timely fashion in mice lacking MLH3 protein, perhaps through repair pathways that differ from those utilized in WT-derived spermatocytes. By contrast, DSBs in *Mlh3^DN/DN^* mutants are either not repaired efficiently or lead to an extended period of DSB induction, resulting in a persistence of RAD51 foci on DSBs through to pachynema.

### *Mlh3^DN/DN^* pachytene spermatocytes show a hyper-accumulation and persistence of BLM

Bloom’s syndrome mutated (BLM) is a mammalian RecQ DNA helicase whose *S. cerevisiae* ortholog, Sgs1, was shown to promote the resolution of complex multi-chromatid joint molecule intermediates that may result from SEI events into both NCOs and COs (Tang et al. 2015; Kaur et al. 2015). During prophase I in WT male spermatocytes, BLM localizes to the chromosomal cores at a high frequency in zygonema and diminishes to a few foci in pachynema (Moens et al. 2000; Walpita et al. 1999; Holloway et al. 2010). Recently, we showed that loss of MLH3 results in up-regulated BLM localization during prophase I, along with persistence of BLM on chromosome cores through late pachynema (Holloway et al. 2010).

To determine if the disruption of the MLH3 endonuclease domain affects the localization of BLM in a similar fashion, to *Mlh3^−/−^*, we stained prophase I chromosome spreads with an antibody against BLM. In zygonema, as previously reported, WT cells show the accumulation of BLM foci on the cores in high numbers, and this frequency is elevated in spermatocytes from both *Mlh3^DN/DN^* and *Mlh3^−/−^* spermatocytes (Figure 3A,B,E,F,I,J,M; p<0.0001 unpaired t-test). This is similar to that reported previously for *Mlh3^−/−^* spermatocytes (Holloway et al. 2010).

**Figure 3.**
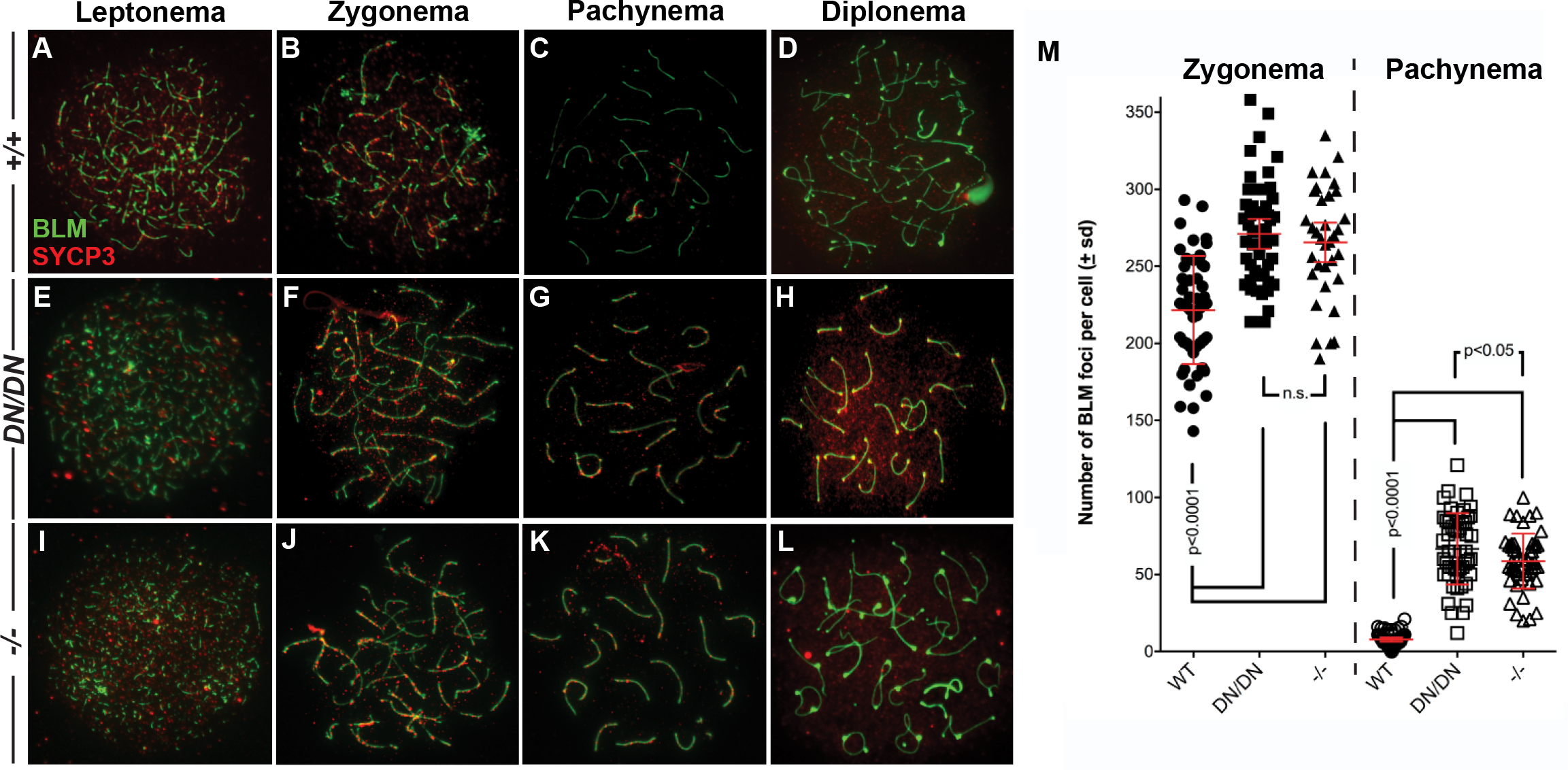
*Mlh3^DN/DN^* spermatocytes show a persistence of BLM in pachynema. Meiotic spreads chromosomes from (A-D) WT, (E-H) *Mlh3^DN/DN^*, and (I-L) *Mlh3^−/−^* males showing the localization of BLM (red) on synaptonemal complex protein SYCP3 (green) throughout the progression of prophase I. (B, F, J, M) In zygonema, *Mlh3^DN/DN^* and *Mlh3^−/−^* cells show significantly more BLM foci localized to the chromosome cores than WT (WT mean ± standard deviation = 221.6 ± 35.1 foci, *Mlh3^DN/DN^* mean ± standard deviation = 271.1 ± 34.2 foci, *Mlh3^−/−^* mean ± standard deviation = 265.5 ± 36.6 foci; p<0.0001 for each using unpaired t-test). (C, D, K, M) In pachynema, BLM is no longer present on the chromosome cores of WT cells whereas *Mlh3^DN/DN^* and *Mlh3^−/−^* cells show hyper-accumulation of BLM on the autosomes and the sex body (WT mean ± standard deviation = 7.8 ± 4.7 foci, *Mlh3^DN/DN^* mean ± standard deviation =66.7 ± 23.2 foci, *Mlh3^−/−^* mean ± standard deviation = 58.7 ± 17.8 foci; p<0.0001 for each using unpaired t-test), which persists into diplonema. At zygonema, the frequency of BLM foci in *Mlh3^DN/DN^* and *Mlh3^−/−^* cells is not statistically different, whereas by pachynema, the number of BLM foci remains significantly elevated in *Mlh3^DN/DN^* spermatocytes, relative to that observed in *Mlh3^−/−^* cells (p<0.05, unpaired t-test). For all chromosome imaging and foci counts, at least 3 mice of each genotype were observed for each staining set.

In early to mid-pachynema, BLM localization on chromosome cores persists in a small percentage of WT spermatocytes, but the number of foci is very much reduced at this stage (Figure 3C,D,M). In contrast, all spermatocytes from *Mlh3^DN/DN^* and *Mlh3^−/−^* spermatocytes show persistent BLM focus localization along chromosome cores (Figure 3G,H,K,L) at a frequency that is elevated above that of WT spermatocytes (Figure 3M, p<0.0001 unpaired t-test). Moreover, the number of BLM foci in *Mlh3^DN/DN^* spermatocytes is significantly elevated relative to that seen in *Mlh3^−/−^* spermatocytes (Figure 3M, p<0.05 unpaired t-test). By late pachynema to early diplonema this difference was even greater, with BLM localization in *Mlh3^DN/DN^* spermatocytes persisting, and being lost in *Mlh3^−/−^* spermatocytes by diplonema (Figure 3D,H,L). Thus, altered MLH3 endonuclease function, like complete loss of MLH3, leads to persistence of BLM helicase on chromosome cores in late prophase I, but at an elevated frequency in *Mlh3^DN/DN^* spermatocytes relative to *Mlh3^−/−^* spermatocytes.

### Increased localization of pro-crossover factor, RNF212, but not MSH4 in pachytene spermatocytes from *Mlh3^−/−^* and *Mlh3^DN/DN^* adult male mice

“Crossover designation” is defined as the process by which class I COs are selected from an excess pool of DSB repair intermediates. In mouse, the 250+ DSBs are processed through zygonema into various repair pathways, and only a subset of these will proceed towards a class I CO fate (Gray and Cohen 2016). These sites become “licensed” for crossing over through the accumulation of the MutS homolog heterodimer, MutSγ (MSH4 and MSH5; (Edelmann et al. 1999; Kneitz et al. 2000)). The MutSγ complex then serves as an early pro-crossover factor by recruiting the MutLγ complex to a select subset of sites and it is these sites that will become “designated” as class I CO events. Notably, all of the 150+ MutSγ sites must be repaired, either as a CO or an NCO, which means that approximately ~125 MutSγ sites must leave the class I CO pathway and undergo repair through an alternate CO pathway or via an NCO pathway, a situation that is unlike that seen in *S. cerevisiae* where the number of MutSγ sites appear to correspond more closely to the number of CO events (Novak et al. 2001).

While the mechanism by which only a subset of MutSγ foci are retained through pachynema remains unclear, studies from a number of groups have implicated the Zip3-like protein, RNF212, in this process (Rao et al., 2017; Reynolds et al., 2013). RNF212 has been shown to co-localize with the majority of MutSγ foci in spermatocytes from WT males and is thought to act as a pro-crossover factor by stabilizing these MutSγ-loaded events (Reynolds et al., 2013). As such, the number of RNF212 foci on chromosome cores is pared down through pachynema in a similar fashion to that of MutSγ (Kolas and Cohen, 2004; Reynolds et al., 2013). Moreover, in mouse mutants that disrupt this paring down process, both RNF212 and MutSγ focus counts remain elevated, but equivalent, throughout prophase I (Holloway et al., 2014; Qiao et al., 2014).

To investigate how loss of MLH3 endonucleolytic function affects this paring down process, we explored RNF212 and MSH4 focus dynamics on chromosome spreads throughout prophase I from WT, *Mlh3^DN/DN^*, and *Mlh3^−/−^* adult male mice (Figure S3). For both RNF212 (Figure S3A-M) and MSH4 (Figure S3N-Z), we find the expected paring down of focus counts from early pachynema (EP) to late pachynema (LP) in spermatocytes from WT, *Mlh3^DN/DN^*, and *Mlh3^−/−^* adult males. In all three cases, RNF212 and MSH4 foci appear on chromosome cores in zygonema (Figure S3B, F, J, O, S, W), persist at high levels in early pachynema (Figure S3C, G, K, P, T, X), and then are reduced to approximately 1-2 foci per chromosome in late pachynema (Figure S3D, H, L, Q, U, Y). Quantification of RNF212 and MSH4 focus numbers in early and late pachytene spermatocytes from WT, *Mlh3^DN/DN^*, and *Mlh3^−/−^* adult male reveals the expected statistically significant decline in these foci EP to LP (Figure S3M,Z; p<0.0001 Mann-Whitney U Test for all). However, the levels of RNF212 foci in both early and late pachynema is significantly higher in spermatocytes from *Mlh3^DN/DN^* and *Mlh3^−/−^* adult males compared to that seen in WT spermatocytes (Figure S3M; p<0.001 Mann-Whitney U test for all). Thus, while the dynamics of RNF212 (high in early and low in late pachynema) are evident in *Mlh3^DN/DN^* and *Mlh3^−/−^* adult males, their focus counts at each of these stages are significantly elevated compared to equivalently-staged WT RNF212 counts. By contrast, at both early and late pachynema, MSH4 counts did not differ between spermatocytes from WT, *Mlh3^DN/DN^* and *Mlh3^−/−^* adult males at both early and late pachynema (Figure S3Z). Thus, mice bearing no MLH3 or catalytically defective MLH3 show a phenotypic divergence in RNF212 and MSH4 focus counts in pachytene spermatocytes.

### Normal localization of Class I Pro-CO factors in pachytene spermatocytes from *Mlh3^DN/DN^* adult male mice

To observe class I CO events in pachynema, we employed two well characterized markers of these sites: the putative ubiquitin E3 ligase, Human Enhancer of Invasion-10 (HEI10), and cyclin-dependent kinase-2 (CDK2) (Ward et al. 2007; Qiao et al. 2014; Ashley et al. 2001). In WT prophase I cells, CDK2 localizes to the telomeres (Figure 4A, yellow arrows) as well as on the chromosome cores (Figure 4A, white arrows) during mid to late pachynema and remains associated with SYCP3 signal through to diplonema (Ashley et al. 2001). The localization of CDK2 along chromosome cores parallels the localization of MLH1 and MLH3, and is associated with nascent class I CO events. In pachytene spermatocyte preparations from *Mlh3^DN/DN^* mice, CDK2 signal remains associated with both the telomeres and chromosome cores at a frequency and intensity that is reminiscent of that seen in WT spermatocyte spreads (Figure 4B). This is in contrast to the situation in spermatocyte preparations from *Mlh3^−/−^* males, in which CDK2 association with the telomere persists, but is lost from nascent CO sites (Figure 4C).

**Figure 4.**
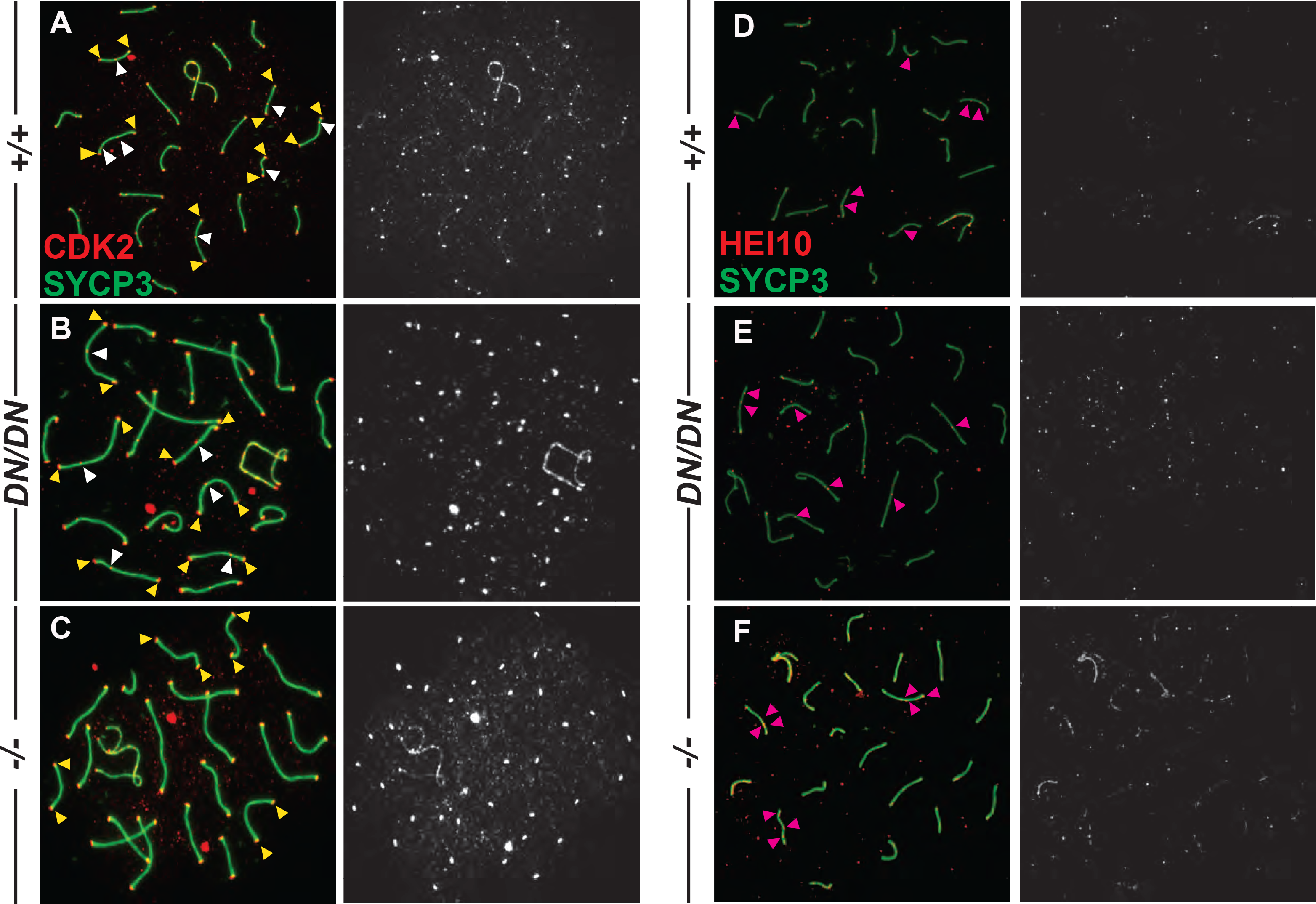
Normal localization of CDK2 and HEI10 to nascent sites of Class I crossovers in *Mlh3^DN/DN^* spermatocytes. (A-C) CDK2 (red) localizes to the synaptonemal complex SYCP3 (green) in WT (A) and *Mlh3^DN/DN^* (B) pachytene spermatocytes, but not in *Mlh3^−/−^* (C) cells, except at the telomeres. Left-hand panels show merged red and green channels, in which white arrows show examples of crossover associated CDK2, while yellow arrows show examples of telomere associated CDK2. Right-hand panels show only the CDK2 signal in white. (D-F) HEI10 (red) co-localizes with the synaptonemal complex SYCP3 (green) in pachytene spermatocytes from WT (D), *Mlh3^DN/DN^* (E), and *Mlh3^−/−^* (F) males. Left-hand panels show merged red and green channels, in which pink arrows show examples of crossover associated HEI10. Right-hand panels show only the HEI10 signal in white. For all chromosome imaging and foci counts, at least 3 mice of each genotype were observed for each staining set.

HEI10 was recently shown to co-localize with MutLγ at sites of class I CO, and its localization is dependent on Cyclin N-terminal Domain-containing-1 (CNTD1)(Holloway et al. 2014; Qiao et al. 2014). HEI10 is thought to play a key role in CO designation/maturation (Qiao et al. 2014). As previously reported for WT cells at pachynema, HEI10 localizes with similar frequency to that of CDK2 and MutLγ (Figure 4D, pink arrows). Similar localization patterns and frequency were observed for *Mlh3^DN/DN^* mice, with a frequency of one to two foci per chromosome (Figure 4E, pink arrows), indicating normal recruitment of HEI10 on pachytene chromosome cores in *Mlh3^DN/DN^* males. This is in contrast to the pattern of HEI10 staining in spermatocytes from *Mlh3^−/−^* mice, where there is an increased accumulation of HEI10 foci (Figure 4F, pink arrows), as previously reported (Qiao et al. 2014). Taken together, these observations demonstrate that loading of HEI10 and CDK2 on class I CO designated sites is affected differently by mutation of *Mlh3:* complete loss of MLH3 results in failure to load CDK2 and hyper-accumulation of HEI10, while altered endonucleolytic activity of MLH3 results in normal loading of both CDK2 and HEI10. Thus, the physical accumulation of MutLγ is required for normal loading of associated pro-crossover maturation factors.

### *Mlh3^DN/DN^* pachytene staged spermatocytes exhibit normal localization of MLH3 and MLH1

MutLγ represents the ultimate marker of DSB repair events that have adopted a class I CO fate, and has been used as a CO proxy marker in many organisms (Lenzi et al. 2005; Auton et al. 2013; Lipkin et al. 2002; Jean et al. 1999). We anticipated that the endonuclease mutation in MLH3 would not affect localization of this complex. In WT spermatocytes, MLH3 localizes on the chromosomes during early pachynema, remaining associated with SYCP3 signal through to diplonema (Figure 5A). In pachytene spermatocyte preparations from *Mlh3^DN/DN^* mice, MLH3 signal remains associated with the autosomal chromosome cores from early pachynema at a focus frequency that is statistically indistinguishable from that of WT cells (Figure 5A-C, p=0.36 by unpaired t-test). MLH3 association with the PAR of the synapsed X and Y chromosomes was similarly unaffected in *Mlh3^DN/DN^* pachytene spermatocytes (data not shown). In addition, the timing of MLH3 appearance, in early pachynema and prior to that of MLH1, was normal in *Mlh3^DN/DN^* pachytene spermatocytes.

**Figure 5.**
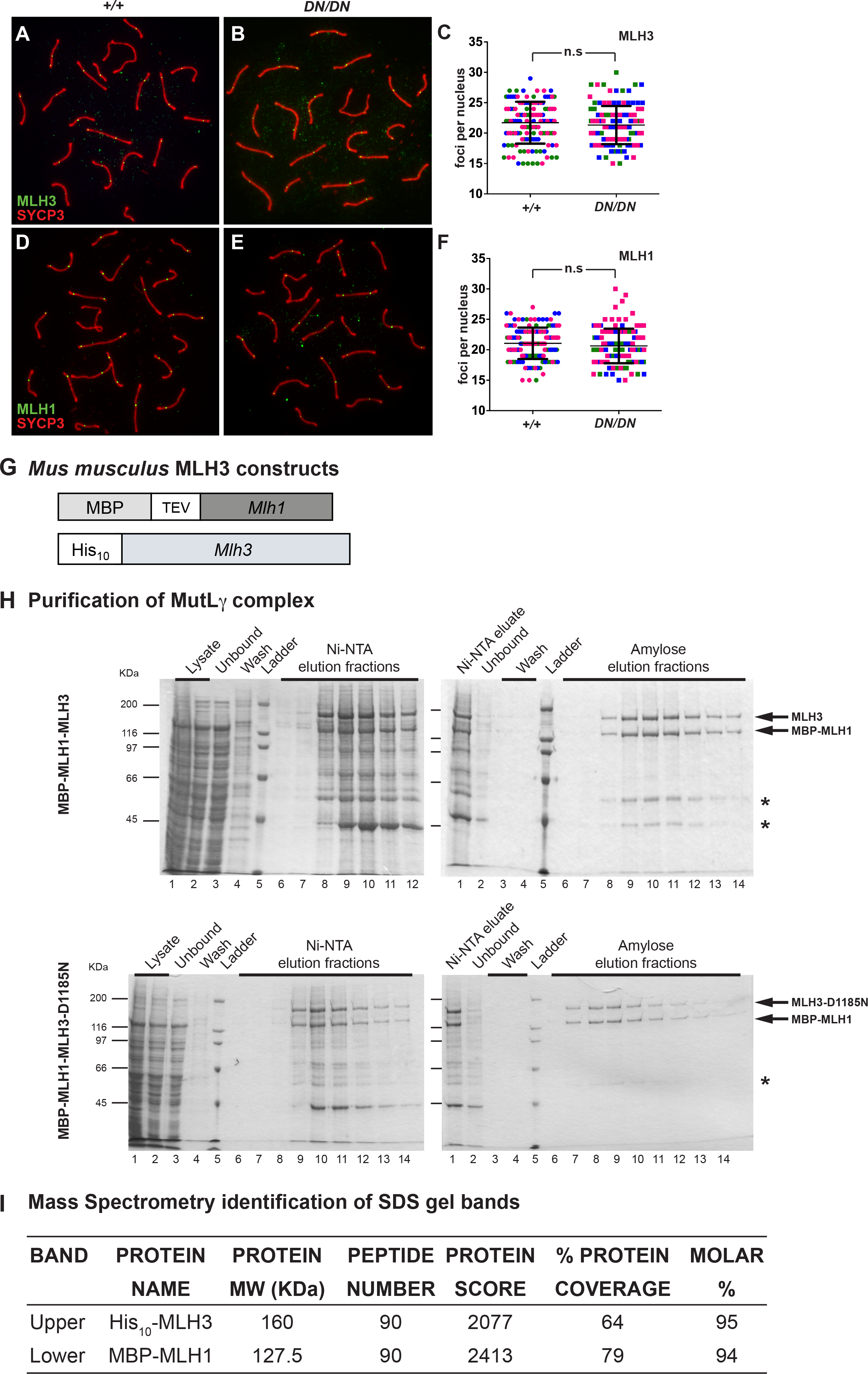
*Mlh3^DN/DN^* pachytene spermatocytes exhibit normal localization of MutLγ while the recombinant MLH3-D1185N protein can form complexes with MLH1. (A-C) MLH3 (green) localizes to SYCP3 (red) in WT and *Mlh3^DN/DN^* pachytene spermatocytes with no statistical difference in the number of MLH3 foci (WT = 21.7 ± 3.4 MLH3 foci n=131, *Mlh3^DN/DN^*= 21.3 ± 3.1 MLH3 foci n=122; p = 0.36 by unpaired t-test). (D-F) MLH1 (green) localizes to SYCP3 (red) in WT and *Mlh3^DN/DN^* pachytene spermatocytes also with no statistical difference in the number of MLH1 foci (WT = 21.1 ± 2.6 MLH1 foci n=144, *Mlh3^DN/DN^* = 20.2 ± 2.8 MLH1 foci n=142; p = 0.2 by unpaired t-test). Different colors in (C) and (F) indicate 3 sets of matched littermates used, each color referring to the counts on a single animal (and color set from the two genotypes being processed simultaneously); n.s = not significant; error bars show standard deviation. (G-I) MLH1-mlh3-D1185N forms a stable heterodimer. (G) Schematic of mouse *Mlh1* and *Mlh3* constructs (see Methods for details). (H) Representative purification of MBP-MLH1-MLH3 (top) and MBP-MLH1-mlh3-D1185N (bottom) using Ni-NTA and amylose resin chromatography as described in the Methods. Fractions were analyzed using SDS-PAGE, stained by Coomassie brilliant blue. MLH1-MLH3 and MLH1-MLH3-D1185N were eluted from amylose in the same fractions. The mass of molecular weight standards is indicated on the left and the expected positions of MBP-MLH1 (127.5 KDa) and His_10_-MLH3 (165 KDa) is indicated in the center. *Likely to be degradation products of MLH1-MLH3. (I) Mass spectrometry analysis of the two major bands in SDS-PAGE detected after amylose chromatography.

Localization of MLH1 was similarly explored in spermatocytes from *Mlh3^DN/DN^* mice. As with MLH3, there was no difference in the timing of MLH1 accumulation on chromosome cores between WT and *Mlh3^DN/DN^* mice (Figure 5D-F). Moreover, when autosomal MLH1 foci were quantified, no statistical difference was observed in MLH1 focus frequency between WT and *Mlh3^DN/DN^* pachytene cells (Figure 5D-F, p= 0.2 by unpaired t-test). These data suggest that disruption of the endonuclease domain of MLH3 does not alter recruitment of MutLγ to chromosomes in pachynema.

### MLH1-MLH3-D1185N forms a stable heterodimer and displays similar chromatographic properties to MLH1-MLH3

Mouse *Mlh1* and *Mlh3* were amplified from cDNA and cloned into pFastBac1 vectors as described in the Methods. The MLH1-MLH3 and MLH1-MLH3-D1185N complexes were expressed from Sf9 cells infected with baculoviruses containing *MBP-Mlh1* and *His_10_-Mlh3* or *His_10_-Mlh3-D1185N* constructs (Figure 5G). Extracts from these cells were applied to a Ni-NTA column. Fractions containing induced proteins were pooled and then applied to an amylose column. Two major bands of molecular weights predicted for an MBP-MLH1-His_10_-MLH3 complex were detected on SDS-PAGE after amylose chromatography (Figure 5H). These bands were further analyzed by mass spectrometry, and the results from this analysis confirmed their identity (Figure 5I). Importantly, MLH1-MLH3 and MLH1-MLH3-D1185N eluted with an apparent 1:1 stoichiometry in both chromatography steps, indicating that the heterodimers were stable, and the protein yields of the two complexes after amylose chromatography were similar (Figure 5H).

### *Mlh3^DN/DN^* diakinesis staged spermatocytes exhibit significantly fewer chiasmata than WT spermatocytes, but elevated chiasmata compared to *Mlh3^−/−^* males

Chiasmata are the physical manifestations of crossing over and, as such, can inform the process of DSB repair via all pathways. Diakinesis-staged spermatocytes from WT and *Mlh3^DN/DN^* males were used to quantify chiasmata. WT cells exhibited a chiasmata frequency of 23.5 ±1.3 per nucleus (Figure 6A, D) whereas *Mlh3^DN/DN^* spermatocytes exhibited a dramatically reduced chiasmata count of 5.2 ± 1.7 chiasmata per nucleus (Figure 6B, D; p <0.0001 by unpaired t-test). Chiasmata counts for *Mlh3^−/−^* males were even more dramatically reduced at 2.8 ± 1.1 chiasmata per nucleus, a value that is significantly lower than both WT and *Mlh3^DN/DN^* spermatocytes (Figure 7C, D; p <0.0001 by unpaired t-test). Thus, complete loss of MLH3 protein leads to the loss of approximately 88% of chiasmata, while loss of endonuclease activity, but retention of MutLγ heterodimer results in only a 78% loss. Thus, the number of residual chiasmata observed in *Mlh3^DN/DN^* spermatocytes is higher than the expected number of chiasmata achieved through the MUS81-EME1-driven class II CO pathway (~2-3, assessed both cytoglogically and genetically; (Svetlanov et al. 2008; Kolas et al. 2005)).

**Figure 6.**
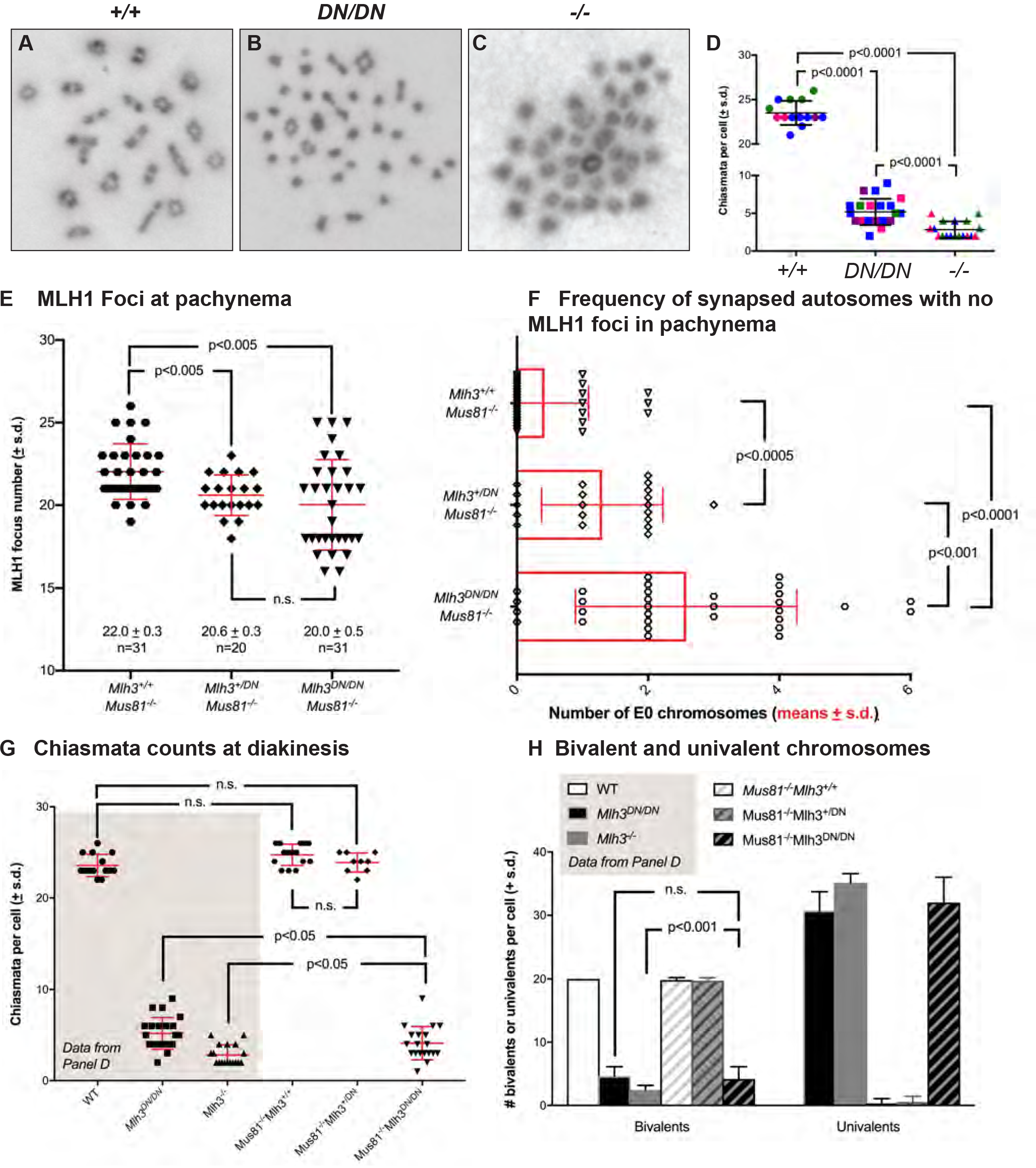
*Mlh3*^*DN/DN*^ diakinesis staged spermatocytes show a reduced number of chiasmata (A-D), while comparison of MLH1 focus frequency (E) and distribution (F) during pachynema, and chiasmata counts (G) and bivalent/univalent frequencies (H) in *Mus81*^*−/−*^*Mlh3*^*+/+*^, *Mus81*^*−/−*^*Mlh3*^*+/DN*^, and *Mus81*^*−/−*^*Mlh3*^*DN/DN*^ males indicates partial involvement of class II CO pathway. (A-C) Diakinesis staged spermatocyte preparations from WT, *Mlh3*^*DN/DN*^, and *Mlh3*^*−/−*^ males stained with Giemsa showing chiasmata formation between homologous chromosomes. (D) *Mlh3^DN/DN^* cells exhibit significantly fewer chiasmata when compared to WT (WT = 23.5 ± 1.3 chiasmata per nucleus, *Mlh3^DN/DN^* = 5.2 ± 1.7; p < 0.0001 by unpaired t-test). *Mlh3*^*−/−*^ cells have significantly fewer chiasmata when compared toWT and *Mlh3^DN/DN^* (*Mlh3*^*−/−*^ = 2.8 ± 1.1 chiasmata per nucleus; p < 0.0001 by unpaired t-test). All values are means + standard deviation. (E-H). We compared MutLγ frequency and distribution during pachynema between mice lacking Mus81 with or without co-incident loss of a endonuclease-intact Mlh3 allele (E, F). *Mus81*^*−/−*^*Mlh3*^*+/+*^ males (E; filled octagons) show elevated MLH1 focus frequency compared to mice bearing one or two copies of the *Mlh3*^*DN*^ allele (filled diamonds and down triangles, respectively). The reduced MLH1 focus count in *Mus81*^−/−^*Mlh3*^*+/DN*^, and *Mus81*^−/−^*Mlh3*^*DN/DN*^ males is statistically significant (p<0.005, unpaired t-test with Welch’s correction). (F) The distribution of MLH1 foci is also disrupted in *Mus81*^−/−^ *Mlh3*^*+/DN*^, and *Mus81*^*−/−*^*Mlh3*^*DN/DN*^ males, with increased numbers of synapsed autosomes showing no MLH1 foci, indicative of non-exchange (E0) chromosome pairs, compared to that seen in *Mus81*^*−/−*^*Mlh3*^*+/+*^ males (statistical significance indicated on the graph; unpaired t-test with Welch’s correction). (G, H) the outcome of class I and class II CO events was assessed by quantifying chiasmata (G) and intact bivalent pairs (H) in diakinesis preparations from mice of all three double mutant genotype combinations, compared to single mutants presented in panel C, above. (G) Chiasmata counts were similar to WT for *Mus81*^−/−^*Mlh3*^*+/+*^ and *Mus81*^−/−^*Mlh3*^*+/DN*^ males (filled octagons and diamonds, respectively), but were statistically significantly lower in *Mus81*^−/−^*Mlh3*^*DN/DN*^ animals (filled down triangles). The frequency of chiasmata in *Mus81*^−/−^*Mlh3*^*DN/DN*^ animals was significantly lower than that observed in *Mlh3^DN/DN^* animals, but significantly higher than that observed in *Mlh3*^−/−^ animals (p<0.05, unpaired t-test with Welch’s correction). However, the number of bivalent structures observed at diakinesis in *Mus81*^−/−^*Mlh3*^*DN/DN*^ animals (panel H) was unchanged from that observed in *Mlh3^DN/DN^* animals. In all cases, two animals were assessed for each genotype.

### Co-incident loss of *Mus81* on the *Mlh3^DN/DN^* background results in altered MutLγ distribution at pachynema and a partial reduction in the residual chiasmata observed in *Mlh3^DN/DN^* single null males

The increased residual chiasmata observed in *Mlh3^DN/DN^* males compared to *Mlh3^−/−^* animals prompted us to ask whether some or all of these crossovers were dependent on the activity of the MUS81-EME1 heterodimer. Previous studies in our lab showed that *Mus81*^−/−^ animals show increased accumulation of MutLγ, resulting in normal chiasmata counts, suggesting that class I CO events are up-regulated in the absence of the class II machinery (Holloway et al. 2008). Co-incident mutation of one or both *Mlh3* alleles to the *Mlh3^DN^* variant on the *Mus81^−/−^* mutant background yielded MLH1 focus counts that were not elevated compared to those observed in *Mus81^−/−^Mlh3^+/+^* mice (Figure 6E, p<0.005 by unpaired t-test with Welch’s correction). Thus, the upregulation of MLH1 foci at pachynema requires both a defective MUS81-EME1 dimer and the presence of only functional MutLγ heterodimer. Interestingly, pachytene spermatocytes from *Mus81^−/−^Mlh3^+/DN^* and *Mus81^−/−^Mlh3^DN/DN^* males show an abnormal distribution of MLH1 foci across all autosomal pairs (Figure 6F). In *Mus81^−/−^Mlh3^+/+^* males, 68% (21/31) cells show no synapsed autosomes without MLH1 foci (so-called “no exchange” or “E0” chromosomes), while in *Mus81^−/−^Mlh3^+/DN^* males, this number dropped to 25% (5/20), with up to 3 E0 chromosomes per cell (Figure 6F). In *Mus81^−/−^Mlh3^DN/DN^* males, the number of synapsed autosomes lacking a MLH1 focus was more significantly increased, with only 13% of cells having no E0 chromosomes, and as many as 6 E0 bivalents being evident (Figure 6F). The increasing frequency of MLH1-devoid autosomes in *Mlh3^+/DN^* and *Mlh3^DN/DN^* males on the *Mus81* null background was statistically significant in all pairwise comparisons (Figure 6F, unpaired t-test with Welch’s correction), perhaps indicating that interference is perturbed in these double mutant animals. In yeast, and also probably in mice, CO interference requires a functional MutSγ complex (Shinohara et al. 2008). However, MutSγ alone is not sufficient to ensure appropriate interference since *Mus81^−/−^* males exhibit disrupted interference despite appropriate MutSγ loading (Holloway et al. 2008).

Assessment of chiasmata counts in single (Figure 6D) and double mutants (Figure 6G) revealed the expected normal chiasmata frequency in spermatocytes from *Mus81^−/−^Mlh3^+/+^* and *Mus81^−/−^Mlh3^+/DN^* males, and the loss of most chiasmata in spermatocytes from *Mus81^−/−^Mlh3^DN/DN^* males. However, whereas the loss of chiasmata structures in *Mlh3^−/−^* and *Mlh3^DN/DN^* single mutants was observed to be 88% and 78%, respectively (Figure 6D), the loss of chiasmata in *Mus81*^−/−^ *Mlh3^DN/DN^* males was 83%. The frequency of chiasmata in cells from these double mutants was statistically different from both *Mlh3* single homozygous mutant animals (p<0.05, unpaired t-test with Welch’s correction). Thus, loss of the class II pathway in addition to the mutation of the endonuclease domain of *Mlh3* only partially reduces residual chiasmata counts to the levels observed in *Mlh3*^−/−^ animals. Interestingly, despite reduced chiasmata, the overall number of bivalent structures observed in diakinesis preparations from *Mus81*^−/−^*Mlh3^DN/DN^* males is the same as that seen in *Mlh3^DN/DN^* males, and is significantly elevated above that seen in *Mlh3*^−/−^ single mutant males (Figure 6H). Taken together, these observations suggest two important features regarding class I/II interactions: (1) that a fully functional class I and class II machinery is required for appropriate distribution of MutLγ foci across the genome; and 2) that MUS81-EME1 activity only partially accounts for the increased residual chiasmata count observed in *Mlh3^DN/DN^* males.

## DISCUSSION

Studies in *S. cerevisiae* and *M. musculus* have implicated MutLγ as the major resolvase of dHJs in the Class I CO pathway (Al-Sweel et al. 2017; Kadyrov et al. 2006; Lipkin et al. 2002; Manhart et al. 2017; Rogacheva et al. 2014; Ranjha et al. 2014; Nishant et al. 2008; Zakharyevich et al. 2012; Edelmann et al. 1996). The current study examines for the first time the importance of an intact endonuclease domain for the proper functioning of MLH3 during prophase I of mammalian meiosis, and is the first exploration of a functional point mutation for MutLγ in the mouse. We generated a mouse lacking a functional endonuclease domain within MLH3, while allowing for heterodimer assembly and near-normal stability of the protein. We found that, as in *Mlh3^−/−^* mutant males, *Mlh3^DN/DN^* males are infertile, exhibit significantly smaller testes than their WT litter mates, and have no epididymal spermatozoa. Beyond this, our data reveal important similarities and differences in the mutant phenotypes to that observed with a nullizygous *Mlh3^−/−^* allele.

We demonstrate that an intact endonuclease domain within MLH3 is not required for DSB or synaptonemal complex formation in early prophase I, similar to that seen in *Mlh3^−/−^* males (Lipkin et al. 2002). However, there are distinct differences in RAD51 accumulation and persistence in spermatocytes from *Mlh3^−/−^* and *Mlh3^DN/DN^* males, suggesting that the effect of MutLγ loss on DSB repair processing is quite different from the presence of a defective MutLγ complex. Most importantly, while RAD51 is recruited in elevated numbers to chromosome cores of *Mlh3^DN/DN^* males, it fails to be cleared effectively in pachynema, perhaps because the defective MutLγ complex blocks subsequent processing of DSB repair intermediates. Intriguingly, the significantly altered RAD51 accumulation in leptonema in both *Mlh3* mutants indicates a role for MutLγ prior to pachynema, far earlier than has been defined thus far. Indeed, analogous early pairing roles for MutLγ have been proposed for *M. Sordaria* and *S. cerevisiae* (Storlazzi et al. 2010; Claeys Bouuaert and Keeney 2017). In yeast, Al-Sweel *et al* constructed whole genome recombination maps for wild-type, endonuclease defective, and null *mlh3* yeast mutants. Both the endonuclease defective and null yeast mutants for *mlh3* showed increases in the number of NCO events, consistent with recombination intermediates being resolved through alternative recombination pathways (Al-Sweel et al. 2017). Thus, in the case of yeast, loss of Mlh3 protein, or the production of an endonuclease defective protein, increases the frequency of other recombination outcomes, most notably including earlier NCO events.

Complete loss of either component of MutLγ eradicates 90-95% of chiasmata, based on the established dogma that class I COs account for the majority, but not all, chiasmata in mammalian meiosis (Baker et al. 1996; Edelmann et al. 1996; Lipkin et al. 2002; Holloway et al. 2008; Svetlanov et al. 2008). By contrast, loss of MUS81, the major class II CO regulator, results in normal chiasmata levels as a result of up-regulation of class I events, as evidenced by a ~10% increase in MutLγ localization during pachynema (Holloway et al. 2008), suggesting that loss of the class II pathway leads to a compensatory increase in class I events. Furthermore, our previous analysis of *Mus81^−/−^Mlh3^−/−^* double mutant animals revealed a very small (~1-2), but consistent, number of residual chiasmata, indicating the existence of other resolvase complexes (Holloway et al. 2008). Taken together, these observations have two important implications for crossing over in the mouse: firstly, additional class I CO events can be achieved through recruitment of additional MutSγ-designated precursor sites in the absence of the class II pathway (and possibly under other circumstances too), and secondly, a few crossovers can be achieved without implementing either MutLγ or MUS81-EME1.

In the current study, we show that spermatocytes from *Mlh3^DN/DN^* males show more residual chiasmata at diakinesis than do *Mlh3^−/−^* mice: approximately 78% of COs are lost in *Mlh3^DN/DN^* spermatocytes, leading to 22% residual chiasmata. This suggests several possibilities: either that the endonucleolytic function of MLH3 does not account for the resolution of all class I COs under normal situations, and/or that other resolvases can be recruited under certain circumstances once MutLγ loads, irrespective of whether this complex is endonucleolytically competent. Alternatively, it is feasible that the point mutation in the endonuclease domain does not completely eliminate endonucleolytic activity in the mouse, resulting in partial class I resolvase activity. We find this latter possibility unlikely due to the severity of the defect in endonucleolytic activity in the *S. cerevisiae* mlh3-D523N protein (Nishant et al. 2008; Rogacheva et al. 2014), but we were unable to test this in the current analysis.

Another explanation for the difference in chiasmata count between *Mlh3^DN/DN^* and *Mlh3^−/−^* mice is that, in the former, some class I CO events can be processed by other CO machinery under conditions of normal accumulation of pro-crossover factors, MutSγ, HEI10 and CDK2, and following processing by DNA helicases such as BLM. In *Mlh3^−/−^* (Holloway et al., 2010) and *Mlh3^DN/DN^* (current work) males, prophase I spermatocytes show increased and persistent accumulation of BLM helicase. A similar increase in BLM localization was also noted in *Mus81^−/−^* spermatocytes (Holloway et al. 2008). In *S. cerevisiae,* loss of class I CO pathway components (for example, in *msh4/5* or *mlh1/3* mutants) is suppressed by mutation of the BLM ortholog, *Sgs1,* highlighting the role of Sgs1 as an anti-crossover factor, and suggesting that the loss of COs in these single MMR mutants is in part due to the activity of Sgs1 (Oh et al. 2007; Jessop et al. 2006). In *msh4/5 sgs1* double mutant strains, the restoration of COs occurs without any concomitant decrease in NCO events, suggesting either that other CO pathways account for the non-class I COs, or that these DSBs are repaired via inter-sister repair processes. In this sense, Sgs1 has been proposed to be master orchestrator of recombination pathway choice (De Muyt et al. 2012), while the Sgs1-Top3-Rmi1 complex as a whole can regulate CO formation both positively and negatively in yeast (Kaur et al. 2015; Tang et al. 2015).

The situation we observe in *Mlh3^−/−^* and *Mlh3^DN/DN^* males with respect to BLM persistence appears to be similar to that seen in yeast. What, then, is the function of BLM at these dHJs that cannot now be processed by MLH3? Previous work in yeast provides a clue to the functional interaction between MutLγ and BLM: crossing over in the class I pathway depends on MutSγ-driven designation on one side of the DSB, in part by suppressing Sgs1, while Sgs1 also imposes an anticrossover bias on the other side of the DSB (Oh et al. 2007). Collectively these two functions promote appropriate CO fates at designated DSB repair sites, preventing the accumulation of unconventional CO/NCO variants through the use of the anti-recombination activity of Sgs1. Extrapolating to the situation in spermatocytes from our two *Mlh3* mutant alleles, we can postulate that the loss of MLH3 protein entirely in *Mlh3^−/−^* males results in a compensatory, but ineffective, increase in BLM that cannot overcome the failure to process class I COs sufficiently (a situation that is different to yeast). In the presence of intact, but catalytically inert MutLγ, on the other hand, the availability of additional BLM can then direct DSB repair in favor of other CO pathways in a similar fashion to the situation in yeast, where the engagement of Sgs1 promotes alternative repair mechanisms, primarily through the recruitment of structure specific nucleases, and the resolution of some dHJs through a class II (or other) CO pathway (De Muyt et al. 2012; Jessop et al. 2006; Jessop and Lichten 2008; Oh et al. 2007; Oh et al. 2008).

While we cannot assess the recruitment of the class II machinery to sites of DSB repair in prophase I in the mouse (due to the lack of available reagents), there is evidence to support the idea that MUS81-EME1 might participate in this CO resolution cross-talk. First, double mutants lacking *Mus81* and bearing a homozygous *Mlh3^DN^* allele show reduced chiasmata relative to *Mlh3^DN/DN^* males, indicating that MUS81 accounts for at least some of the increase in chiasmata above that of *Mlh3^−/−^* males. Second, normal accumulation of MSH4 in *Mlh3^DN/DN^* spermatocytes recruits normal levels of pro-crossover factors, HEI10 and CDK2, despite reduced RNF212 levels, suggesting that other CO resolution pathways might be engaged at the expense of class I pathways.

We provide evidence herein that COs are achieved in *Mlh3^DN/DN^* spermatocytes in a manner that is partially dependent on the MUS81-EME1 endonuclease. There are other resolvase complexes that might also generate COs in mammalian meiosis, and these cannot be ruled out in the current analysis. Indeed, our previous analysis of *Mus81^−/−^Mlh3^−/−^* males indicated the existence of additional CO events that were independent of the class I and class II pathways (Holloway et al. 2008). Additional resolvases in yeast include SLX1-SLX4 and YEN1/GEN1(Zakharyevich et al. 2012; Arter et al. 2018; Muñoz-Galván et al. 2012). Indeed, the persistence of DSB repair intermediates into pachynema, along with the upregulated and persistent BLM might suggest that the defective MutLγ complex prevents accumulation of other such resolvase complexes. This might, in turn delay CO maturation until later in prophase I when, for example, GEN1 can be invoked to resolve the CO (García-Luis and Machín 2014). Thus, we propose that the timing of MutLγ activity, and its clearance from nascent COs is an important factor in the recruitment of alternative CO processing machineries, but in a manner that is not dependent on its endonuclease activity.

## Materials and Methods

### Generation of mice and genotyping

PL253 targeting vector containing the *Mlh3-D1185N* point mutation in the potential endonuclease domain and a loxP-neo-loxP cassette in intron 5-6 of *Mlh3* was incorporated into an embryonic stem cell line. *Mlh3^DN^* transgenic mice were crossed with a *Spo11-Cre* mouse line to remove the *neo* cassette (Lyndaker et al., 2013), and then maintained on an inbred background through backcrossing on to the C57Bl/6J line (Jackson Laboratory, Bar Harbor, ME). Genotyping of WT, *Mlh3^+/DN^*, and *Mlh3^DN/DN^* mice was performed using the following PCR primer pairs: forward (5’-AAGCCAAGTCTGCATGAGTA-3’) and reverse (5’-TAAATGTGCCACTGACTAAAT-3’) followed by a restriction enzyme digestion with *Sau96I* (New England Biolabs) at 37ºC for 2-3 hours, which results in 439-bp and 263-bp fragments from the WT allele and a 702-bp fragment from the mutant allele. Fertility tests were performed by breeding *Mlh3^DN/DN^* adult males with WT females. At least 3 males of each genotyped were evaluated. Presence of a copulation plug the following morning counted as a successful mating event. Pregnancy was confirmed by gentle palpation of the abdomen after gestation day 11 or on delivery date of litters. Mice were housed and utilized under the guidance and approval of the Cornell University Institutional Animal Care and Use Committee.

### Histology

Testes from adult mice were fixed in Bouin’s solution overnight at room temperature and then washed 3 x 10 min with 70% ethanol at room temperature with agitation. Fixed and paraffin-embedded testes were section at 5 μm. H&E staining was performed on Bouin’s fixed testes using standard methods. At least 6 males of each genotyped were evaluated.

### Sperm counts

Caudal epididymides were removed from adult males and placed in pre-warmed 1X PBS containing 4% bovine serum albumin. Sperm were released into solution by squeezing epididymis with tweezers and incubated for 20 min at 32ºC/5% CO_2_. After incubation, 20 μL of sperm suspension was re-suspended in 480 μL of 10% formalin. Sperm counts were performed with a hemocytometer. At least 10 males of each genotype were evaluated for sperm counts and testis weights.

### Prophase I chromosome analysis and immunofluorescence

Prophase I chromosome spreads from adult testes were prepared as previously described (Kolas et al., 2005). For all experiments, at least 6 males of each genotyped were evaluated. Chromosome slides were then washed in 0.4% Kodak Photo-Flo 200/1X PBS for 2 x 5 min, 0.4% Kodak Photo-Flo 200/dH_2_O for 2 x 5 min, then air-dried for approximately 10 min and stored in −80°C or used immediately for staining. Primary antibodies used were: anti-γH2AX (Millipore, NY, #05-636 1:10,000), anti-SYCP3 (Abcam, MA, #97672, 1:5000), anti-SYCP1 (Abcam, MA, #15087, 1:1000), anti-RAD51 (Calbiochem, #PC130, 1:500), anti-BLM (generous gift from Dr. Ramundo Freire; 1:100;), anti-CDK2 (Santa Cruz, TX, sc-163; 1:250), anti-MLH3 ((Holloway et al., 2014); 1:1000;), anti-RNF212 (generous gift from Dr. Neil Hunter), anti-MSH4 (Abcam, MA, #58666), anti-HEI10 (Anti-CCNB1IP1, Abcam, MA # 71977) and anti-MLH1 (BD Biosciences Pharmingen, CA, #550838, 1:100). Secondary antibodies used were: goat anti-mouse Alexa Fluor 488 (#62-6511), goat anti-mouse Alexa Fluor 555 (#A-10521), goat anti-rabbit Alexa Fluor 488 (#65-6111), goat anti-rabbit Alexa Fluor 555 (#A-10520; all Invitrogen, 1:2000).

### Spermatocyte diakinesis spread preparations to observe chiasmata

Diakinesis chromosome spreads were prepared as previously described (Evans et al., 1964; Uroz et al., 2008) with slight modifications (Holloway et al., 2010). Slides were stained with 10% Giemsa for 10 mins, washed, air-dried and mounted with Permount.

### Image acquisition

All chromosome spread slides were visualized using the Zeiss Imager Z1 microscope (Carl Zeiss, Inc.). Images were captured with a high-resolution microscopy camera AxioCam MRM (Carl Zeiss, Inc.) and processed with ZEN Software (version 2.0.0.0; Carl Zeiss, Inc.).

### Cloning mouse *Mlh1* and *Mlh3*

cDNA was synthesized from total testis RNA from wildtype C57B/6J adult males using the SuperScript III Reverse Transcriptase Kit from ThermoFisher. *Mlh1* and *Mlh3* open reading frames were PCR amplified from cDNA using Expand High Fidelity DNA polymerase using primer pairs AO3365 (5’GCTAGCAGCTGATGCATATGGCGTTTGTAGCAGGAG) and AO3366 (5’TACCGCATGCTATGCATTAACACCGCTCAAAGACTTTG) for *Mlh1*, and AO3367 (5’ACGTCGACGAGCTCATATGCATCACCATCACCATCACCATCACCATCACATCAGGTGTCTATCAGATGAC) and AO3368 (5’CGAAAGCGGCCGCGATCATGGAGGCTCACAAGG) for *His_10_*-*Mlh3*. Each fragment was cloned into the *Spe*1 site of pFastBac1 (ThermoFisher) using Gibson assembly PCR (NEB) to create pEAE393 (*Mlh1*) and pEAE397 (*His_10_-Mlh3*). Constructs were verified by DNA sequencing with NCBI reference sequences NM_026810.2 and NM_175337.2 for *Mlh1* and *Mlh3,* respectively. These constructs were then modified as follows:

1. The MBP–TEV sequence was inserted at the N-terminus of *Mlh1* in pEAE393 to create pEAE395.
2. The *mlh3-D1185N* mutation was introduced into pEAE397 by Quick Change (Agilent) to create pEAE413.

### Chromatography analysis of the mouse MLH1-MLH3-D1185N heterodimer from Baculovirus-infected Sf9 cells

*Sf9* cells were transfected with pEAE397 (*His_10_-Mlh3*), pEAE413 (*His_10_-Mlh3-D1185N*) and pEAE395 (*MBP-Mlh1*) using the Bac-to-Bac baculovirus infection system (Invitrogen). Fresh *Sf9* cells were co-infected with both viruses (containing *Mlh1* and *Mlh3* or *Mlh3-D1185N*). Cells were harvested 60 hours post infection, washed with phosphate buffered saline, and kept at −80°C until use.

Cell pellets from 250 ml of cells as thawed, resuspended in 60 ml hypotonic lysis buffer (20 mM HEPES-KOH pH 7.5, 5 mM NaCl, 1 mM MgCl_2_, 1 mM PMSF and EDTA free protease inhibitor mixture from Roche and Thermo Scientific) and incubated for 15 min on ice. The suspension was adjusted to 250 mM NaCl, 15 mM imidazole, 10% glycerol, 2 mM ß-mercaptoethanol (BME), and clarified by centrifugation at 17,000 g for 20 min at 4°C. The supernatant was mixed with 6 ml of 50% nickel-nitrolotriaceticacid-agarose (Ni-NTA) resin and allowed to bind for 2 hours or overnight followed by centrifugation to remove the unbound fraction. The resin was packed onto a column and washed with 7-10 column volumes of wash buffer (50 mM HEPES-KOH pH 7.5, 250 mM NaCl, 40 mM imidazole, 10% glycerol, 2 mM BME, 1 mM PMSF). Protein was eluted with 15 ml of 300 mM imidazole in 50 mM HEPES-KOH pH 7.5, 250 mM NaCl, 40 mM imidazole, 10% glycerol, 2 mM BME and 1 mM PMSF. Elution fractions containing MLH1-MLH3, determined by SDS-PAGE, were pooled and loaded onto 1 ml 100% amylose resin (NEB). The resin was washed with 10 column volumes of wash buffer (50 mM HEPES-KOH pH 7.5, 250 mM NaCl, 10% glycerol, 2 mM BME, 1 mM PMSF) and eluted with 6 ml wash buffer containing 10 mM maltose. Fractions containing MLH1-MLH3 were pooled and aliquots were flash frozen and stored in −80°C. The protein yield, following amylose chromatography, was similar for wild-type and mutant complexes (approximately 120-150 μg per 250 ml cells).

It is important to note that we were unable to detect a specific endonuclease activity for the mouse MBP-MLH1-MLH3 complex, suggesting that the MBP tag interferes with MLH1-MLH3 functions. We were unable to test this directly because, despite numerous attempts, we were unable to efficiently remove the MBP tag from MLH1 by treating MBP-MLH1-MLH3 with TEV protease.

### Mass-spectrometry of MLH1 and MLH3 bands from SDS-PAGE

SDS-PAGE bands following amylose chromatography predicted to contain MBP-MLH1 and His_10_-MLH3 were excised and analyzed by the Cornell University Proteomics facility using a Thermo LTQ Orbitrap Velos Mass Spectrometer.

### Statistical methods and analysis

The majority of comparisons involved with unpaired parametric t-test with Welch’s correction or nonparametric Mann-Whitney U-test, depending on the data distribution. All statistical analysis was performed with GraphPad Prism Version 7.00 for Mac, Graphpad Software, La Jolla California USA, www.graphpad.com. P-values less than 0.05 were considered statistically significant.

## Acknowledgements

The authors of this work would like to thank Dr. Andrew Miller for necropsy analysis of WT, *Mlh3^DN/DN^* and *Mlh3^−/−^* mice (data not included), Dr. Ramundo Freire for providing anti-BLM antibody, and Dr. Neil Hunter for providing anti-RNF212 antibody. We are grateful to members of the Cohen lab for their critical evaluation of this work. This work was supported by the National Institutes of Health (NIH) grant R01HD041012 to P.E. Cohen, and by graduate student funding through The Training Grant in Reproductive Genomics (PI Mark S. Roberson, T32HD052471) for M. Toledo. V. Raghavan and E. Alani were supported by funding to E. Alani from The NIH (grant GM53085). The authors declare no competing financial interests.

## Author contribution

Melissa Toledo performed experiments, curated data, supervised undergraduates, and wrote the original draft; Xianfei Sun performed experiments, generated the mutant allele, and conceptualized the project; Miguel A. Brieño-Enríquez performed experiments, curated data, and reviewed/edited the manuscript; Vandana Raghavan performed experiments and analyzed data; Stephen Gray performed experiments, curated data, and reviewed/edited the manuscript; Jeffrey Pea performed experiments and analyzed data; Anita Venkatesh performed experiments; Lekha Patel performed experiments; Peter L. Borst performed experiments; Eric Alani curated data, conceptualized experiments, analyzed data, and reviewed/edited the manuscript; Paula E. Cohen acquired funding, supervised the experiments, curated data, conceptualized experiments, wrote the manuscript, and reviewed/edited the manuscript.

---

**Figure S1. Amino acid sequence of MLH3 endonuclease domain and its conservation across species.** (A) Amino acid protein sequence of the *M. musculus, H. sapiens, S. cerevisiae, and A. thaliana* MLH3 endonuclease domain, DQHA(X)_2_E(X)_4_E, shows the conservation of this domain across these species. Asterisk refers to the conserved aspartic acid (D) that was targeted for a point mutation and converted to asparagine (N) to generate the *Mlh3-DN* mouse. (B) *Mus musculus* MLH3 is composed of a 1411 amino acid long sequence that results in an ~158 kDa sized protein (UniProt, 2015). MLH3 contains a globular N-terminal domain (NTD; light gray) and C-terminal domain (CTD; light blue) connected by a flexible linker arm (white). The NTD (light gray) contains ATP binding motifs (dark gray) that are conserved across species. The CTD consists of the MLH1 interacting domain (light blue) and the conserved endonuclease motif (dark blue). The aspartic acid (D) in the conserved endonuclease motif in mouse, DQHAAHERIRLE, was converted to an asparagine (N; red) at amino acid site 1185.

**Figure S2. DSB formation and signaling as well as synapsis are normal in *Mlh3^DN/DN^* spermatocytes throughout prophase I.** (A-O) *Mlh3^DN/DN^* prophase I cells exhibit normal DSB formation and signaling as observed by γH2AX staining (green) on synaptonemal complex protein SYCP3 (red) as compared to WT and *Mlh3^−/−^* cells. Images show abundant γH2AX signal in leptonema, following by diminished signal in zygonema, with the absence of signal in pachynema and diplonema, except at the sex body, due to MSCI. (E, J, O) Over exposure of the γH2AX signal results in γH2AX foci or flares on the autosomes in WT, *Mlh3^DN/DN^*, as well as *Mlh3^−/−^* cells (white arrows), suggesting that this signal may not be representative of true un-repaired DSBs. (P-W) *Mlh3^DN/DN^* prophase I cells have normal synapsis as observed by the localization of synaptonemal complex protein SYCP1 (green) and SYCP3 (red) on the chromosomes when compared to WT. SYCP3 forms as short patches along the chromosomes in leptonema, extending into filaments in zygonema along with the appearance of SYCP1, full synapsis with the co-localization of SYCP1 and SYCP3 are observed in pachynema, followed by desynapsis in diplonema with the degradation of SYCP1.

**Figure S3. Normal localization of crossover designation factor MSH4 in *Mlh3^DN/DN^* and *Mlh3^−/−^* spermatocytes.** (A-M) RNF212 (green) localization on chromosome cores stained with antibodies against SYCP3 (red) throughout prophase I (leptonema, zygonema, early pachynema, and late pachynema). Diplonema and diakinesis are not shown. Representative images of spermatocytes from WT (A-D), *Mlh3^DN/DN^* (E-H), and *Mlh3^−/−^* (I-L) adult males. Panel M shows the quantitation of RNF212 foci in all three genotypes at early (EP) and late pachynema (LP). RNF212 accumulates on chromosome cores at zygonema in high numbers and these foci diminish gradually through pachynema with only one or two foci remaining in late pachynema. (N-Z) MSH4 (green) localization on chromosome cores stained with antibodies against SYCP3 (red) throughout prophase I (leptonema, zygonema, early pachynema, and late pachynema). Diplonema and diakinesis are not shown. Representative images of spermatocytes from WT (M-P), *Mlh3^DN/DN^* (Q-T), and *Mlh3^−/−^* (U-X) adult males. Panel Z shows the quantitation of MSH4 foci in all three genotypes at early (EP) and late pachynema (LP). A similar pattern of MSH4 foci accumulation and loss is observed for MSH4 as for RNF212: accumulation of high numbers of foci in zygonema, diminishing to one or two foci per chromosome in late pachynema. In all cases, statistical analyses was performed using unpaired t-test with Welch’s correction (p values provided in graphs). For all chromosome imaging and foci counts, at least three mice of each genotype were observed for each staining set.

