## Supplementary figures and images for "A mutation in the endonuclease domain of mouse MLH3 reveals novel roles for MutLγ during crossover formation in meiotic prophase I"

### Supplemental Figure S1

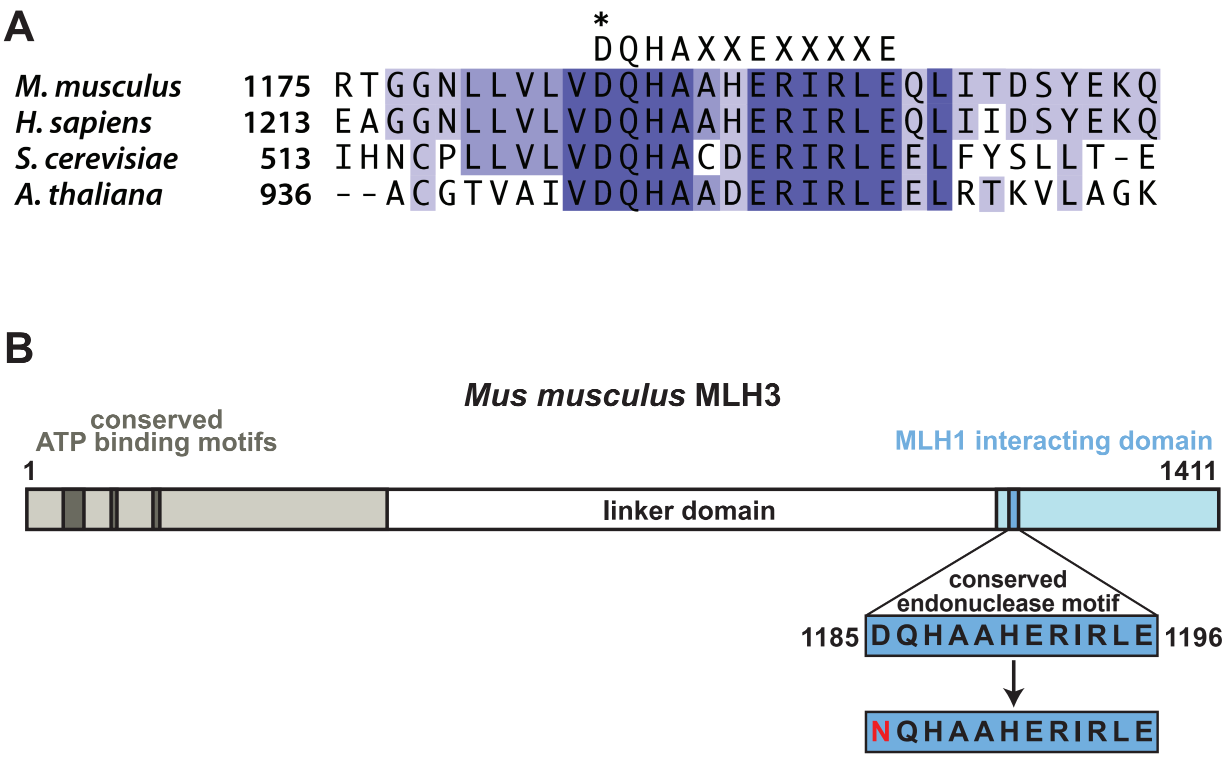

### Supplemental Figure S2

Toledo *et al:* Figure S2.

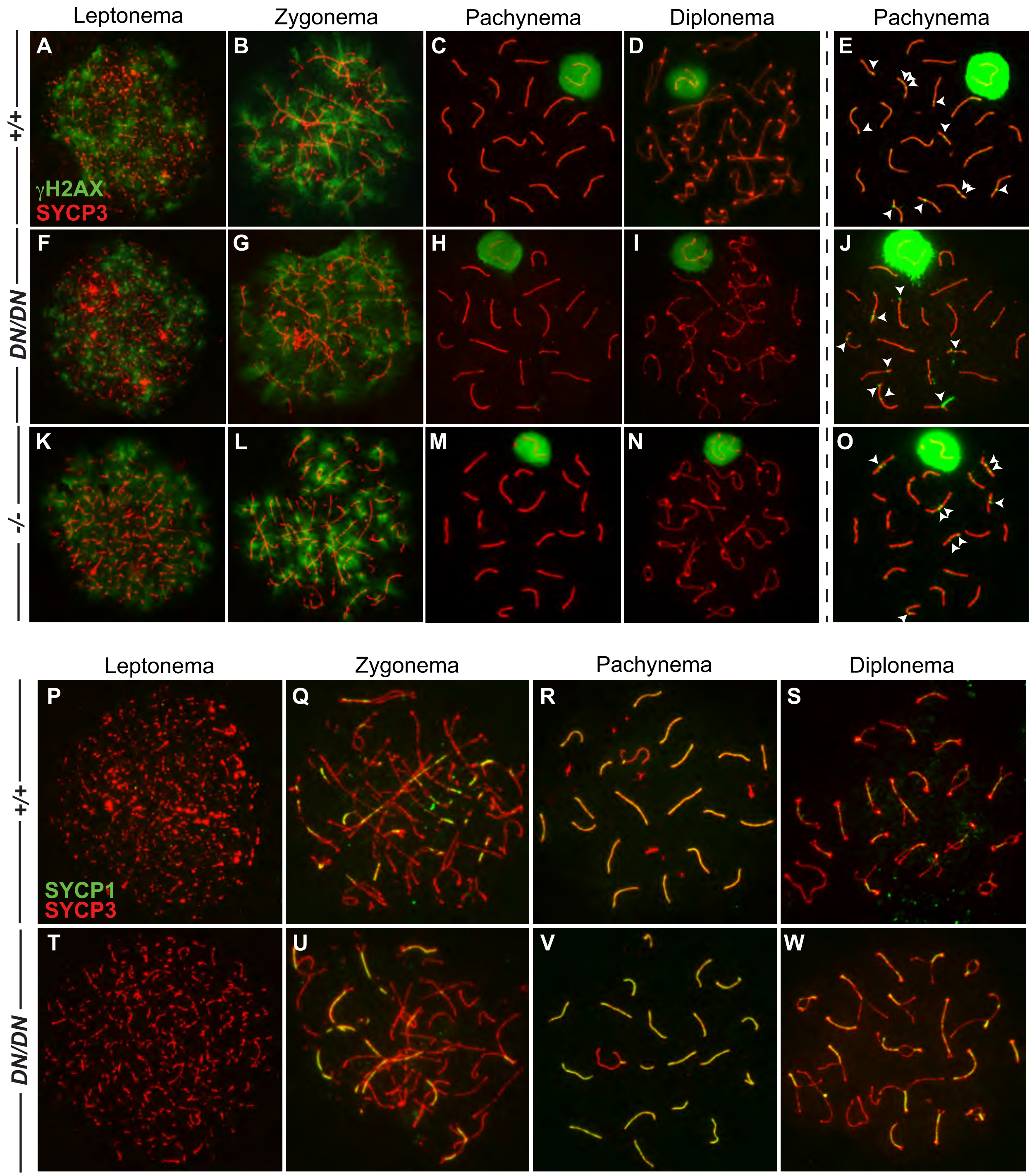

### Supplemental Figure S3

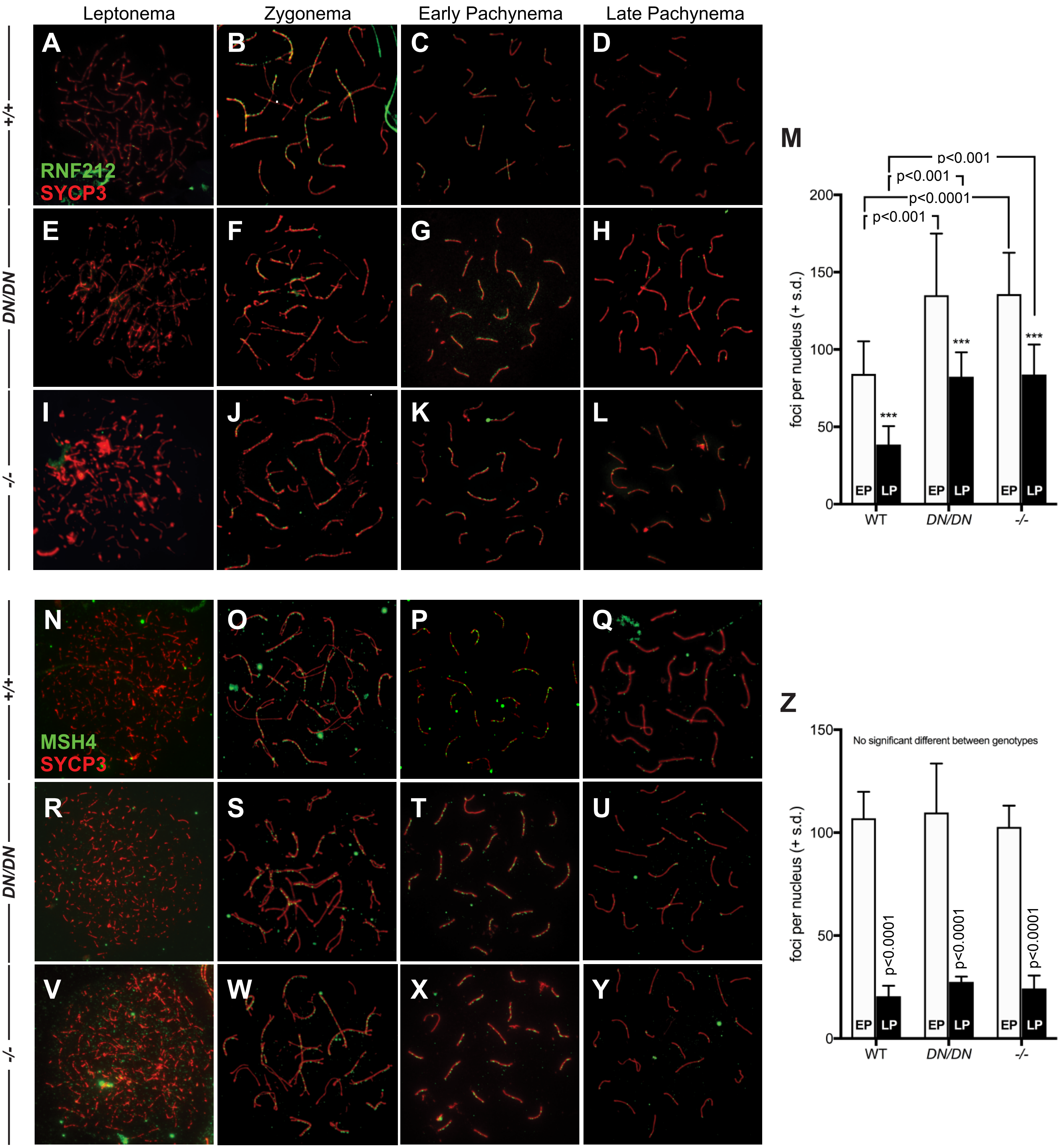
